# Functional crosstalk between phosphorylation and disease-causing mutations in the cardiac sodium channel Na_v_1.5

**DOI:** 10.1101/2020.12.11.417683

**Authors:** Iacopo Galleano, Hendrik Harms, Koushik Choudhury, Keith Khoo, Lucie Delemotte, Stephan Alexander Pless

**Affiliations:** Department of Drug Design and Pharmacology, University of Copenhagen, Copenhagen, 2100 Denmark; Science for Life Laboratory, Department of Applied Physics, KTH Royal Institute of Technology, Solna, SE-171 65 Sweden

**Keywords:** post-translational modifications, protein engineering, cardiac arrhythmias, anti-arrhythmic drugs, sodium channel inactivation

## Abstract

The voltage-gated sodium channel Na_v_1.5 initiates the cardiac action potential. Alterations of its activation and inactivation properties due to mutations can cause severe, life-threatening arrhythmias. Yet despite intensive research efforts, many functional aspects of this cardiac channel remain poorly understood. For instance, Na_v_1.5 undergoes extensive post-translational modification *in vivo*, but the functional significance of these modifications is largely unexplored, especially under pathological conditions. This is because most conventional approaches are unable to insert metabolically stable post-translational modification mimics, thus preventing a precise elucidation of the contribution by these modifications to channel function. Here, we overcome this limitation by using protein semi-synthesis of Na_v_1.5 in live cells and carry out complementary molecular dynamics simulations. We introduce metabolically stable phosphorylation mimics on both WT and two pathogenic long-QT mutant channel backgrounds and decipher functional and pharmacological effects with unique precision. We elucidate the mechanism by which phosphorylation of Y1495 impairs steady-state inactivation in WT Na_v_1.5. Surprisingly, we find that while the Q1476R patient mutation does not affect inactivation on its own, it enhances the impairment of steady-state inactivation caused by phosphorylation of Y1495 through enhanced unbinding of the inactivation particle. We also show that both phosphorylation and patient mutations can impact Na_v_1.5 sensitivity towards the clinically used anti-arrhythmic drugs quinidine and ranolazine, but not flecainide. The data highlight that functional effects of Na_v_1.5 phosphorylation can be dramatically amplified by patient mutations. Our work is thus likely to have implications for the interpretation of mutational phenotypes and the design of future drug regimens.

**Significance statement:** The cardiac sodium channel (Na_v_1.5) is crucial for generating a regular heartbeat. It is thus not surprising that mutations in its sequence have been linked to life-threatening arrhythmias. Interestingly, Na_v_1.5 activity can also be altered by posttranslational modifications, such as tyrosine phosphorylation. Our combination of protein engineering and molecular modeling studies has revealed that the detrimental effect of a long QT3 patient mutation is only exposed when a proximal tyrosine is phosphorylated. This suggests a dynamic crosstalk between the genetic mutation and a neighboring phosphorylation, a phenomenon that could be important in other classes of proteins. Additionally, we show that phosphorylation can affect the channel’s sensitivity towards clinically-relevant drugs, a finding that may prove important when devising patient-specific treatment plans.

## Introduction

The cardiac action potential is elicited by the temporally precisely orchestrated activity of voltage-gated sodium, potassium and calcium channels. The voltage-gated sodium ion channel Na_v_1.5, for which more than 500 potentially pathologically relevant point mutations have been reported (1), is responsible for the initial fast depolarization observed in cardiac action potentials. Many of these mutations are the known or suspected cause of aberrant action potentials that can result in life threatening conditions like Brugada and long QT3 syndromes (BS and LQT3, respectively) (2). The most common non-invasive treatment for such conditions includes administration of anti-arrhythmic drugs (AADs), such as quinidine, flecainide or ranolazine (3).

Na_v_1.5 is a 2016-amino acid membrane protein with four homologous but distinct domains (DI-IV) (4). Each domain contains six transmembrane segments (S1-S6). The first four segments (S1-S4) form the so called ‘voltage sensing domain’, in which the positively charged residues of S4 are responsible for the voltage sensitivity of the channel. Conversely, the last two segments (S5-S6), and the P-loop that connects them, form the pore module, including the selectivity filter. Upon depolarization, the S4 segments undergo an upward movement, which is translated to conformational changes that open the channel gate. This channel opening is primarily mediated by S4 helices in DI-DIII, while the slower upward movement of DIV causes the channel to inactivate (5–8). The latter is caused by the IFM tripeptide motif in the DIII-DIV linker binding to its receptor site adjacent to DIV S6 (9–13). Subthreshold depolarizations can cause the channel to undergo steady-state inactivation (SSI) without prior opening. The voltage dependence of SSI is an important determinant of channel availability *in vivo* and even small alterations can give rise to arrhythmic phenotypes. Similarly, incomplete inactivation can result in a potentially pathogenic standing current referred to as late current (14).

Intriguingly, many of the known or suspected disease-causing mutations occur in the cytosolic interdomain linkers and the loops between individual transmembrane segments (1, 11, 15–17). These loops and linkers are not only hotspots for disease causing mutations, but also contain numerous posttranslational modifications (PTMs), such as phosphorylation, methylation and acetylation (18). Phosphorylation stoichiometries are typically variable (19), and previous work has shown that phosphorylation levels can increase substantially in cardiac ion channels such as Na_v_1.5 and K_v_7.1 upon e.g. β-adrenergic stimulation (20). Consequently, mutations at PTM sites or dysregulation of PTM levels play important roles in disease states (18, 21). Yet despite firm evidence for their presence and relevance *in vivo*, it has remained challenging to directly assess the functional effects of PTMs *in vitro*: conventional mutagenesis can be used to introduce non-modifiable side chains to prevent modification, but mimicking PTMs using naturally occurring side chains is often flawed, especially with mimicking phosphorylation of tyrosine (22). Similarly, overexpression of regulatory enzymes, e.g. kinases or phosphatases, lacks selectivity toward the protein of interest. Recently, we overcame some of these limitations by using tandem protein-trans splicing (tPTS) to generate semi-synthetic membrane proteins containing stable PTM mimetics (23).

Here, we employ a combination of tPTS, electrophysiology and molecular dynamics (MD) simulations to investigate if the functional effects caused by phosphorylation of Na_v_1.5 Y1495 (10 mV right-shift in the V_1/2_ of SSI) (23, 24) are altered by clinically relevant mutations in its proximity and if they affect the pharmacological profile of AADs in WT and mutant channels. Our data show that in contrast to the pathogenic ΔK1500 mutant (25), the disease-causing Q1476R mutation (26) has no significant impact on SSI *per se*. However, when the Q1476R mutation is combined with phosphorylation at the nearby Y1495, this leads to a dramatically right-shifted SSI (20 mV) and a prominent late current. Our MD simulations attribute these shifts in SSI to less stable binding of the IFM inactivation particle in channels with phosphorylation at 1495, in particular when combined with the Q1476R mutation. Finally, we show that Na_v_1.5 sensitivity towards two class I AADs and ranolazine is altered by phosphorylation and the tested patient mutations.

## Results

### Experimental strategy

Heterologous protein expression does not allow precise control over the extent of post-translational modification. In the case of Na_v_1.5 Y1495, while a conventional mutagenesis approach can prevent phosphorylation with a tyrosine-to-phenylalanine mutation, it is not possible to control the degree of phosphorylation on the native tyrosine. We overcome this obstacle using a recently developed semi-synthetic approach which allows for the insertion of synthetic peptides carrying single or multiple PTMs or PTM mimics into ion channels (23). In this approach, the channel is divided into three fragments: N- and C-terminal fragments (N^REC^ and C^REC^) corresponding to partial channel fragments that are recombinantly expressed in *Xenopus leavis* oocytes, and a synthetic peptide (P^SYN^) containing the site of interest, which is injected into the oocyte cytosol (Fig. 1A and Fig. S1). Covalent linkage of the three fragments is mediated by two orthogonal split inteins: *Cfa*DnaE (split intein A) (27) and *Ssp*DnaB^M86^ (split inteins B) (28). Upon assembly, the respective split inteins spontaneously and covalently link the attached channel fragments via native chemical ligation, a process also termed tandem protein trans-splicing (tPTS) (29) (Fig. 1A, Fig. S1 and Fig. S2).

**Fig. 1.**
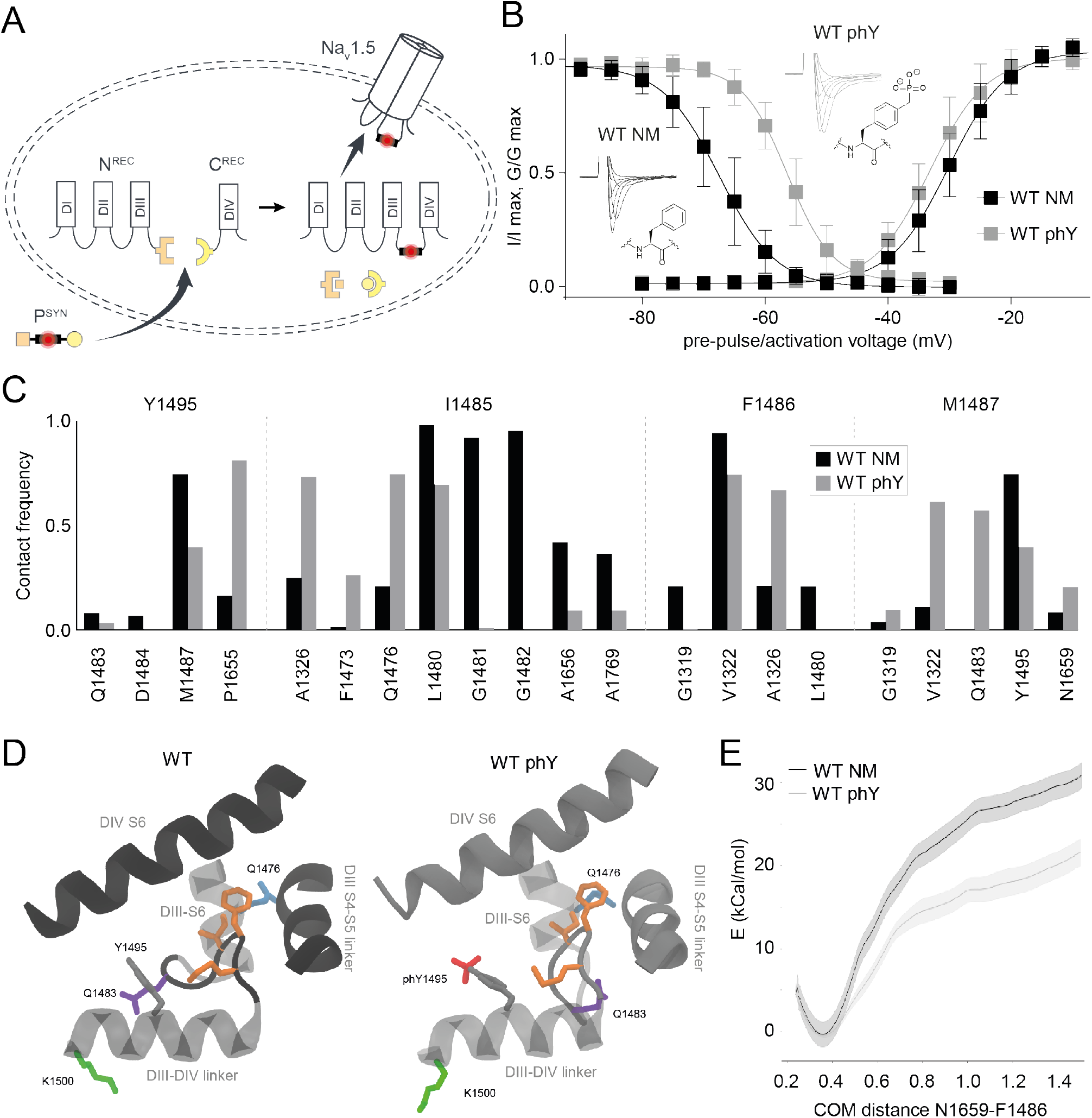
Phosphorylation of Y1495 destabilizes docking of IFM motif into its receptor site. (A) Schematic of tPTS used to generate Na_v_1.5 channels that are non-modifiable (WT NM) or phosphorylated (WT phY) at the Y1495 position. Amino acids (aa) 1-101 of *Cfa*DnaE (orange) are merged to the C-terminus of channel fragment corresponding to Na_v_1.5 aa 1-1471 for heterologous expression as the N^REC^ construct. The P^SYN^ sequence corresponds to Na_v_1.5 aa 1472-1502 and is linked to the C-terminal part of *Cfa*DnaE (aa 102-137, orange) at its N-terminus and the N-terminal part of *Ssp*DnaB^M86^ (aa 1-11, yellow) at its C-terminus. The corresponding C-terminal part of *Ssp*DnaB^M86^ (aa 12-154, yellow), is expressed as a fusion construct at the N-terminus of protein fragment C (Na_v_1.5 aa 1503-2016) to form the C^REC^ construct. (B) SSI (left) and activation (right) curves of WT NM and WT phY constructs, including example traces and chemical structure of the amino acid present in position 1495. Data shown as mean ± standard deviation; n = 6-9. (C) Contact frequency of Y1495 and IFM particle residues with neighboring residues. Phosphorylation reduces the contact between Y1495 and M1487 of IFM while it causes the IFM particle to increase contacts with DIII-S6 and DIII-S4-S5 linker (D) Conformation of IFM particle in its binding site after 200 ns of MD simulation. Phosphorylation of Y1495 moves the loop region of the DIII-DIV linker containing the Q1483 residue outward and the IFM particle closer to the DIII-S6. IFM side chains highlighted in orange. (E) Free energy profile of IFM unbinding, using the distance between the center of mass of N1659 in the DIV-S5 helix and F1486 in the IFM particle as a reaction coordinate. WT phY causes a decrease in the binding energy of IFM binding.

This approach allows us to insert P^SYN^ variants either containing a phenylalanine (which cannot be post-translationally modified, hence designated non-modifiable, NM) or a phospho<u>n</u>ylated tyrosine (30) in position 1495 (phY). The latter is a hydrolytically stable phosphorylation mimic with near identical charge and steric properties (31). We will thus refer to effects of phosphorylation throughout the manuscript whenever we introduce phY. To increase splicing efficiency, we introduced the N1472C mutation at the first position of P^SYN^, which results in a right-shift in SSI compared to WT channels ((23) and Table 1). This is consistent with the notion that the highly similar N1472S variant has previously been reported as a putative long QT-associated mutation (32).

**Table 1:** Electrophysiological characterization. Values for half-maximal activation (V_50_ Activation, in millivolts) and SSI (V_50_ SSI, in millivolts) are listed, along with late currents observed 400 ms after the peak current elicited by a voltage step to −30 mV (I _late_ (% of I _peak_)) and the time constant for inactivation in response to a voltage step to −20 mV (τ _inactivation_, in milliseconds) as well as the time constant for recovery from inactivation in response to two voltage steps to − 20 mV with varying recovery intervals at − 80 mV in between (τ _recovery_, in milliseconds). Data provided as mean ± standard deviation; the number of replicates for each value is shown in brackets; nd, not determined.

| Construct | V<sub>50</sub> Activation<br>(mV) | V<sub>50</sub> SSI<br>(mV) | I<sub>late</sub><br>(% of I<sub>peak</sub>) | τ<sub>inactivation</sub><br>(ms) | τ<sub>recovery</sub><br>(ms) |
| --- | --- | --- | --- | --- | --- |
| WT | - 37.3 ± 2.3<br>(9) | - 76.7 ± 1.9<br>(9) | nd | nd | nd |
| WTrec | - 34.4 ± 3.1<br>(7) | - 67.7 ± 3.5<br>(8) | nd | nd | nd |
| WT NM | - 30.1 ± 2.9<br>(7) | - 67.5 ± 3.6<br>(7) | - 4.6 ± 2.7<br>(6) | 1.3 ± 0.3<br>(7) | 20.9 ± 4.3<br>(7) |
| WT phY | - 32.8 ± 3.2<br>(7) | - 56.4 ± 2.2<br>(6) | - 2.4 ± 3.2<br>(7) | 1.7 ± 0.3<br>(7) | 9.3 ± 1.5<br>(5) |
| Q1476R NM | - 38.3 ± 2.5<br>(7) | - 68.1 ± 2.5<br>(7) | - 2.2 ± 1.9<br>(10) | 1.8 ± 0.2<br>(6) | 14.8 ± 1.5<br>(6) |
| Q1476R phY | - 33.2 ± 6.7<br>(6) | - 47.8 ± 5.0<br>(6) | 10.7 ± 4.5<br>(5) | 2.9 ± 0.7<br>(8) | 6.7 ± 1.4<br>(7) |
| ΔK1500 NM | - 29.1 ± 2.6<br>(7) | - 71.3 ± 3.4<br>(6) | 14.5 ± 2.8<br>(6) | 3.0 ± 0.5<br>(6) | nd |
| ΔK1500 phY | - 33.0 ± 3.5<br>(6) | - 59.4 ± 3.2<br>(7) | 9.9 ± 5.4<br>(10) | 3.1 ± 0.8<br>(6) | nd |
| Y1495E | - 30.0 ± 2.0<br>(10) | - 55.8 ± 2.1<br>(11) | nd | nd | nd |
| Q1476R+Y1495E | - 30.9 ± 1.5<br>(6) | - 50.7 ± 2.3<br>(10) | nd | nd | nd |

### Phosphorylation of Y1495 alters interaction between IFM tripeptide and receptor site

Injection of oocytes expressing N and C fragments with NM peptide (WT NM) results in activation and SSI parameters similar to those obtained when all three channel fragments are expressed recombinantly (WT^rec^) (Fig. S3 and Table 1). However, no voltage-dependent currents were observed within similar incubations times when only N and C fragments were injected, suggesting that the channel fragments alone were unable to non-covalently assemble (23). Consistent with previous work, phospho<u>n</u>ylation at Y1495 (WT phY) results in a roughly 10 mV right-shift in the SSI curve, while retaining WT-like values for half-maximal activation (Fig. 1B and Table 1) (23, 33). The time course of inactivation remained unchanged and no late current was observed, while the time course of recovery from inactivation of WT phY was significantly accelerated relative to the one of WT NM (Fig. S4 and Table 1).

To elucidate the molecular basis for these experimental observations, we turned to molecular dynamics simulations of the recently determined cryo-EM structure of Na_v_1.5 in which the IFM particle is docked to a site between the DIII S4-S5 linker and DIV S6 in an intermediate inactivated state (11). Since SSI is attributed to this specific interaction (Fig. 1D), we focused on differences between WT NM and WT phY in this region. Strikingly, 200 ns MD simulations reveal that phosphorylation causes a loss in contact of Y1495 with M1487 of the IFM particle. In WT phY, a contact between I1485 of IFM and F1473 and Q1476 (in DIII-S6) appears (I1485, Fig. 1C). Interactions involving F1486, on the other hand, displays no substantial differences between these two conditions, except for the formation of contacts with A1326 upon phosphorylation (F1486, Fig. 1C). The phosphate group of WT phY mediates a downward movement of Q1483 and moves D1484 away from DIV-S6 (Y1495, Fig. 1C,D). Thus, upon phosphorylation of Y1495, the phosphate group moves towards the binding pocket of the IFM particle (Fig. 1D), which in turn pushes the IFM particle towards DIII-S6, making contacts with Q1476. This causes the N-terminal part of the DIII-DIV linker (Q1483, D1484) to move out and downwards, away from the IFM binding pocket (Fig. 1D).

Umbrella sampling simulations were carried out to evaluate the binding free energy of the IFM particle to its docking site, using the distance between the center of mass of N1659 in DIV-S5 helix and F1486 in the IFM particle as a reaction coordinate (Fig. 1E). This specific reaction coordinate was chosen because N1659 forms a hydrogen bond with the carbonyl group of F1486 in the docked state, as suggested by Jiang et. al. (11). As expected from the observations from regular MD simulations, phosphorylation of Y1495 indeed causes the binding free energy to decrease, suggesting that the unbinding of the IFM particle is eased by phosphorylation. This is in good agreement with the right-shift in SSI and the accelerated recovery from inactivation observed experimentally.

Together, the combined experimental and computational data support the notion that phosphorylation at Y1495 destabilizes the interaction of the IFM motif with its receptor site. Specifically, phosphorylation of Y1495 moves it away from the DIV-S6 and towards the DIII-S6 and DIII-S4-S5 linker, thereby right-shifting Na_v_1.5 SSI and accelerating recovery from inactivation.

### Q1476R pathogenicity likely arises from phosphorylation at Y1495, not the mutation per se

The Q1476R mutation in the DIII-DIV linker had previously been suggested to be the cause of LQT3 in a patient family (26). Specifically, heterologous expression in mammalian cells had demonstrated that the mutant channels displayed a 6.5 mV right-shift in SSI compared to WT, along with the emergence of a late current (26). Given the close physical proximity of Q1476 and Y1495 with the IFM inactivation particle, and the increased contact frequency of I1485 and Q1476 observed in WT phY, we sought to test if phosphorylation of Y1495 would have a distinct effect on channel function in the presence of the Q1476R mutation. To this end, we synthesized P^SYN^ variants containing the Q1476R mutation on either the Y1495F (Q1476R NM) or the Y1495 phY background (Q1476R phY). Insertion of either peptide variant resulted in robust voltage-gated currents within 12 hrs after injection (Fig. 2A). To our surprise, introducing the Q1476R mutation on the non-phosphorylated background did not shift the SSI voltage dependence or elicit a late current (Fig. 2B,C and Table 1). By contrast, when we introduced the Q1476R mutation in the presence of a phosphorylated tyrosine at 1495, we observed a drastic right-shift in SSI compared to Q1476R NM (ΔSSI ~20 mV, Table 1), significantly beyond what we observed on the WT background (ΔSSI ~11 mV). We further show that phosphorylation on the mutant (but not the WT) background also significantly slowed the rate of channel inactivation and resulted in a pronounced late current after 400 ms (Fig. 2A,C and Table 1). The only parameter that was significantly affected by Q1476R alone (Q1476R NM), was the voltage required for half-maximal activation. This was significantly left-shifted compared to WT NM, but indistinguishable for Q1476R phY and WT phY (Fig. 2B,C and Table 1).

**Fig. 2.**
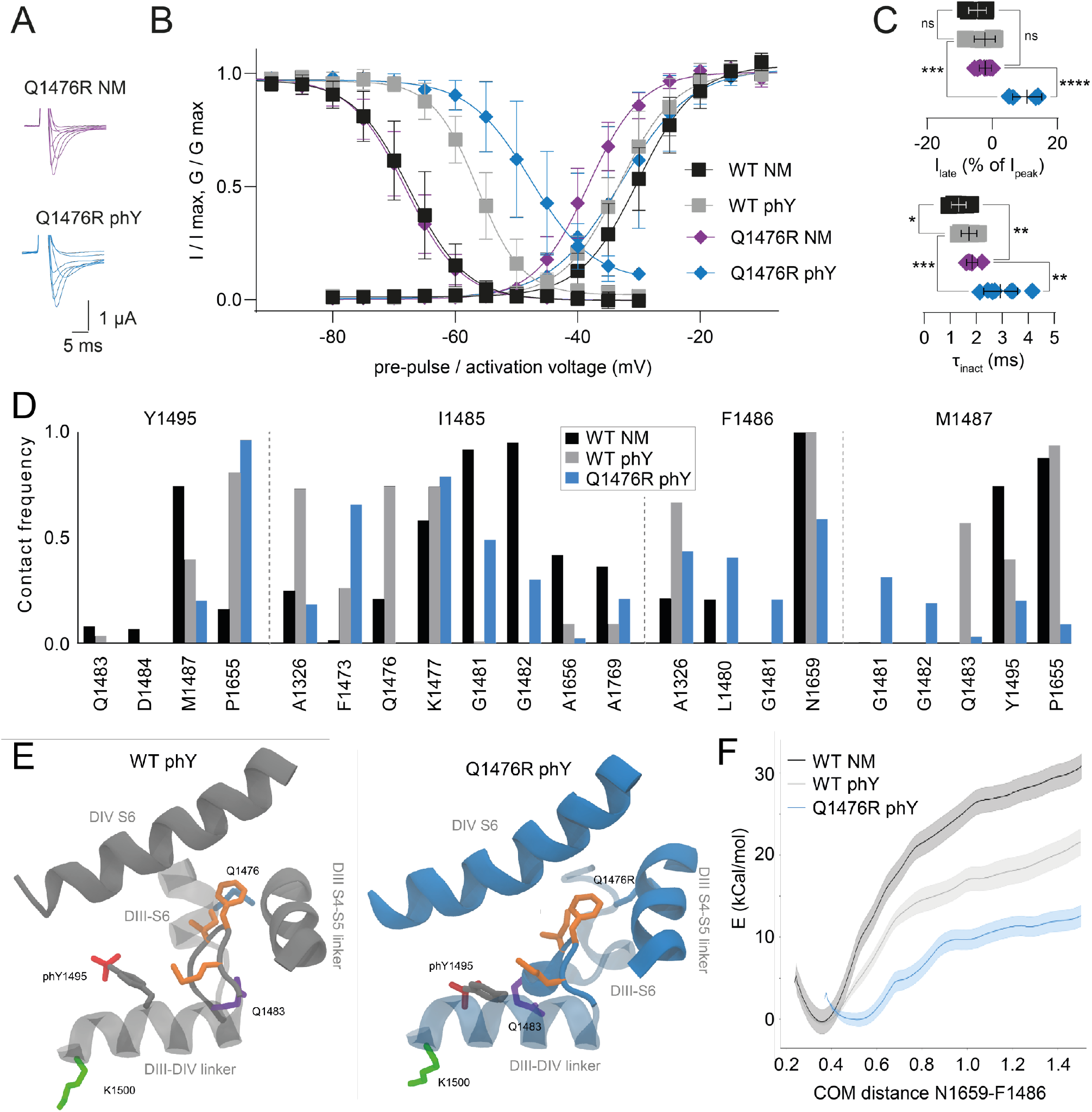
The Q1476R mutation results in a dramatically shifted SSI. (A) Representative current traces for Q1476R NM (purple) and phY (blue) constructs. (B) SSI (left) and activation (right) curves of indicated constructs. (C) Late currents (upper panel) and inactivation rates (lower panel) for indicated constructs. Data shown as mean ± standard deviation in (B) and (C); n = 5-10, data was compared using unpaired two-tailed student’s t-test, ns (not significant) p > 0.05, * p ≤ 0.05, ** p ≤ 0.01, *** p ≤ 0.001, **** p < 0.0001. Note that the precise measurements of late currents in *Xenopus laevis* oocytes are hampered by slow-onset, voltage-dependent endogenous currents. Therefore, our late current measurements can only serve as an estimate. (D) Contact frequency of Y1495 and IFM particle with neighboring residues. (E) Conformation of IFM particle in its binding site after 200 ns of MD simulation. Q1476R phY leads to a break in the DIII-S6 helix, leading to the destabilization of IFM binding. IFM side chains highlighted in orange. (F) Free energy profile of IFM unbinding, using the distance between the center of mass of N1659 in DIV-S5 helix and F1486 in the IFM particle as a reaction coordinate. Q1476R phY results in a further decrease of binding energy of the IFM particle compared to WT phY and an increased equilibrium distance between F1486 and its docking site due to the break in DIII-S6 induced by the Q1476R mutation.

These data suggest that the phosphorylation-induced destabilization of the IFM particle in its receptor site was starkly exacerbated in the presence of the pathogenic Q1476R mutation. This idea is further supported by our finding that recovery from inactivation was accelerated for Q1476R phY compared for WT phY (Fig S4 and Table 1). Next, we sought to assess if full-length Na_v_1.5 containing a conventional glutamic acid as a mimic of phosphotyrosine in position 1495 on either the WT or the Q1476R background would yield comparable results (Y1495E and Q1476R/Y1495E, respectively). Although we observed overall similar trends, both Y1495E and Q1476R/Y1495E resulted in even more pronounced shifts in SSI compared to the semi-synthetic WT NM, WT phY and Q1476R phY constructs (Fig. S3 and Table 1). This underscores the advantages of inserting phosphorylation mimics with near identical charge and steric properties via our semi-synthetic approach, rather than using conventional mutagenesis.

To rationalize the molecular basis for the observed effect caused by Q1476R, we conducted 200 ns long MD simulations of Na_v_1.5 with both the WT phY and the Q1476R phY systems. Strikingly, the Q1476R mutation causes a break in the DIII-S6 helix, allowing its C-terminal end to move into the IFM binding site. As a consequence, increased contact frequency was observed between G1481/G1482 and the IFM motif in the Q1476R phY system compared to the WT phY system (Fig. 2D,E). The Q1476R mutation also leads to a loss of contact of position 1476 itself to I1485, suggestive of the importance of this contact for stabilizing the IFM motif in the WT phY system. We also observe a decrease in contact frequency between F1486 of IFM and N1659 of DIV-S6 and the formation of new contacts with L1480 and G1481 (Fig. 2D,E). Finally, M1487 reduces its interaction with P1655, Q1483 and phY in position 1495, while increasing its contact with G1481 and G1482. Our umbrella sampling results also indicate that the combination of Q1476R and phY further decreases the binding free energy of the IFM motif to its binding site and leads to a larger equilibrium distance between the F1486 and its docking site, in line with expectations from regular MD simulations (Fig. 2F).

Overall, we show that the effects of Y1495 phosphorylation on SSI are strongly enhanced in the presence of the Q1476R patient mutation, while the latter has little effect in the absence of the Y1495 phY modification. In WT phY, we observe increased contacts of IFM with DIII-S6, in particular through contacts between I1485 and Q1476. Thus, mutating Q1476 to R appears to further destabilize the IFM interactions with their docking site in a phosphorylated state. Together, this implies that the pathogenicity of the Q1476R mutation does not arise from the amino acid change *per se*, but rather the functional effects conferred by a nearby phosphorylation.

### Phosphorylation at Y1495 has similar effects on ΔK1500 and WT channels

Next, we set out to assess if other patient mutants in close proximity to Y1495 would show a similar differential modulation by phosphorylation. Specifically, we chose a deletion mutant, ΔK1500, that had previously been associated with LQT3, BS and conduction system disease (25). To this end, we synthesized P^SYN^ variants containing the ΔK1500 deletion mutation on either the Y1495F (ΔK1500 NM) or the Y1495 phY background (ΔK1500 phY). With both, we observed robust voltage-gated currents (Fig. 3A). Interestingly, the midpoints of voltage dependence for activation and SSI were very similar to those observed for the respective WT constructs (compare ΔK1500 NM and WT NM; ΔK1500 phY and WT phY; Fig. 3B/C and Table 1). In other words, phosphorylation at position 1495 in the presence of ΔK1500 results in a ~12 mV right-shift in SSI, which is nearly identical to the shift observed on the WT background. However, the ΔK1500 mutation does reduce the slope of the SSI curve and results in a decreased rate of inactivation and a distinct late current (Fig. 3A-C and Table 1), even in the absence of phosphorylation.

**Fig. 3.**
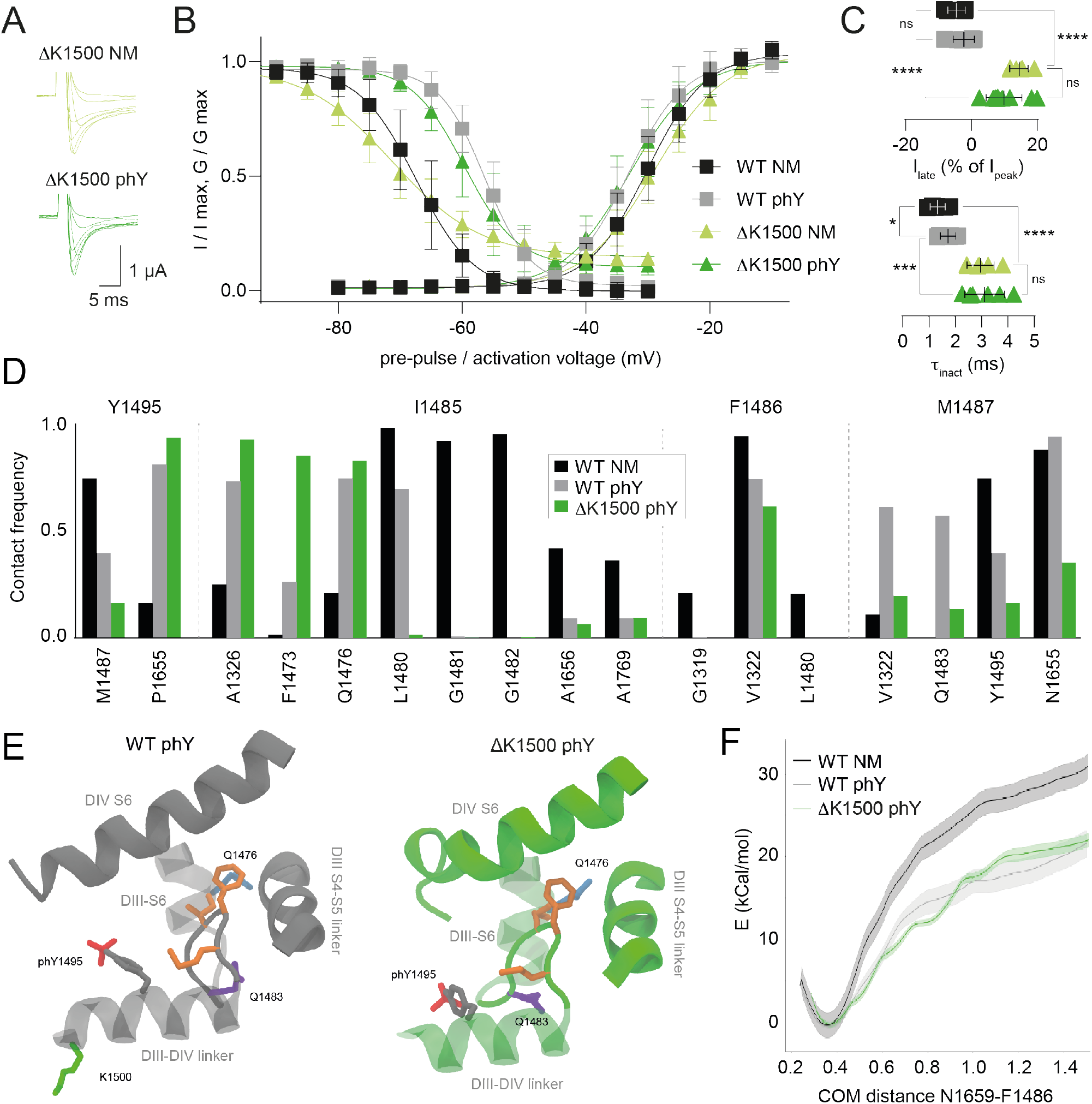
Phosphorylation-induced SSI shift is similar in ΔK1500 and WT. (A) Representative current traces for ΔK1500 NM (light green) and ΔK1500 phY (dark green) constructs. (B) SSI (left) and activation (right) curves of indicated constructs. (C) Late currents (upper panel) and inactivation rates (lower panel) for indicated constructs. Data shown as mean ± standard deviation in (B) and (C); n = 6-10, data was compared using unpaired two-tailed student’s t-test, ns (not significant) p > 0.05, * p ≤ 0.05, *** p ≤ 0.001, **** p < 0.0001. (D) Contact frequency of Y1495 and IFM particle with neighboring residues. (E) Conformation of the IFM particle in its binding site after 200 ns of MD simulation. I1485 increases its contacts with DIII-S6 residues. IFM side chains highlighted in orange. (F) Free energy profile of IFM unbinding, using the distance between the center of mass of N1659 in the DIV-S5 helix and F1486 in the IFM particle as a reaction coordinate. The binding energy of the IFM particle for the ΔK1500 phY system is similar to that of WT phY.

Our observation that phosphorylation affects the inactivation properties of ΔK1500 in a manner similar to WT is further corroborated by our computational studies. We find the binding free energy of the IFM particle to its binding site to indeed be indistinguishable from that of WT phY (Fig. 3F), despite a slightly altered interaction profile between the IFM particle and its binding site (Fig. 3D,E).

Together, we show that the ΔK1500 mutation *per se* slows the rate of inactivation and results in a substantial late current, thus likely explaining its pathogenicity. However, and in contrast to the Q1476R mutation, the functional effects elicited by phosphorylation at Y1495 are virtually identical to those observed with WT channels.

### Differential effects of phosphorylation and patient mutations on Na_v_1.5 pharmacology

To test if the above observations have consequences for Na_v_1.5 pharmacology, we next investigated if phosphorylation of Y1495 alone or in combination with the Q1476R or the ΔK1500 patient mutations would affect the channel’s sensitivity towards clinically used AADs. First, we tested the Class Ia AAD quinidine (Fig. 4A), which displays pronounced use-dependent inhibition of Na_v_1.5. Inhibition by quinidine was assessed using a 20 Hz stimulation protocol and yielded an IC_50_ of 86 ± 83 μM for WT NM (Fig. 4A,C and Table 2). By contrast, phosphorylation of Y1495 increased the IC_50_ to 159 ± 47 μM (WT phY; Fig. 4A,C and Table 2). Introduction of the Q1476R mutation led to a further decrease in apparent affinity, but this value was no longer affected by phosphorylation at Y1495 (206 ± 196 μM for Q1476R NM vs of 195 ± 96 μM for Q1476R phY; Fig. 4A,C and Table 2). Similarly, on the ΔK1500 mutant background, quinidine inhibition was virtually independent of phosphorylation at Y1495, with IC_50_ values of 63 ± 55 μM (ΔK1500 NM) and 43 ± 5 μM (ΔK1500 phY), respectively (Fig. 4A,C and Table 2).

**Table 2:** Pharmacological sensitivity towards clinically relevant AADs. Half-maximal inhibitory concentrations (IC_50_) for quinidine, flecainide and ranolazine. Values are provided as mean ± standard deviation, the number of replicates for each value is shown in brackets.

| **Construct** | **IC<sub>50</sub> quinidine**<br>**(μM)** | **IC<sub>50</sub> flecainide**<br>**(μM)** | **IC<sub>50</sub> flecainide**<br>**(μM)** |
| --- | --- | --- | --- |
| **WT NM** | 86 ± 83<br>(6) | 23 ± 12<br>(8) | 60 ± 28<br>(8) |
| **WT phY** | 159 ± 47<br>(9) | 33 ± 20<br>(6) | 156 ± 68<br>(6) |
| **Q1476R NM** | 206 ± 196<br>(7) | 23 ± 5<br>(5) | 80 ± 35<br>(7) |
| **Q1476R phY** | 195 ± 96<br>(7) | 21 ± 9<br>(5) | 130 ± 68<br>(6) |
| **ΔK1500 NM** | 63 ± 55<br>(6) | 23 ± 8<br>(9) | 57 ± 25<br>(6) |
| **ΔK1500 phY** | 43 ± 5<br>(7) | 16 ± 6<br>(6) | 146 ± 67<br>(7) |

**Fig. 4.**
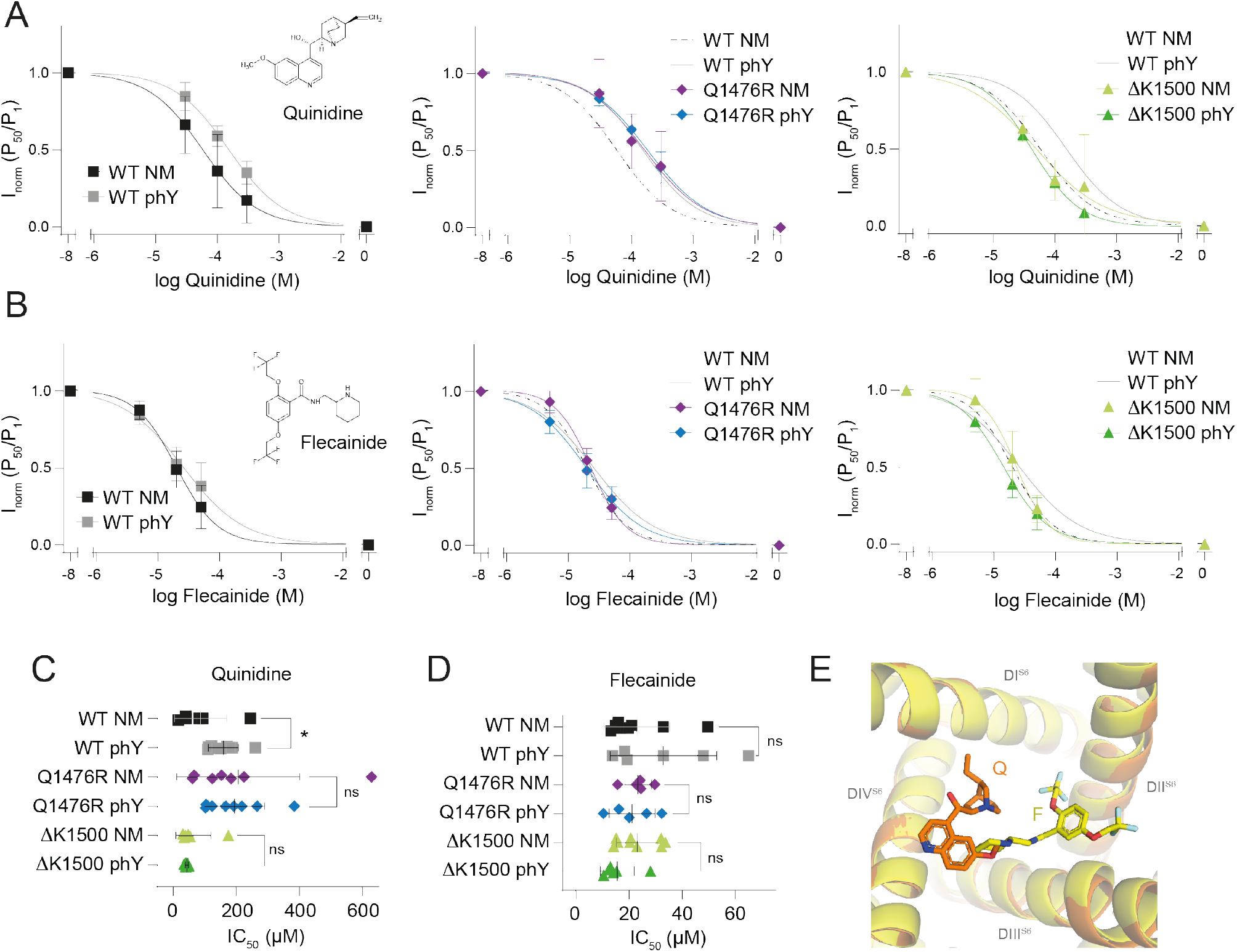
Phosphorylation and disease mutations can affect pharmacological sensitivity of Na_v_1.5. (A) and (B) Concentration response curves of WT, Q1476R, and ΔK1500 constructs in response to a 20 Hz pulse train stimulation in presence of antiarrhythmic drugs quinidine (A) or flecainide (B), respectively (see structures in left panels). P_50_/P_1_ values were normalized to range from 0 to 1. (C) IC_50_ values obtained for quinidine data shown in (A). The IC_50_ is significantly increased by phosphorylation only in WT, but not in Q1476R or ΔK1500 constructs. (D) IC_50_ values obtained for flecainide data shown in (B). IC_50_ values are not significantly altered by phosphorylation in any of the constructs. ns (not significant) p ≥ 0.05, * p < 0.05. Data shown as mean ± standard deviation; n = 5-9. (E) Overlay of flecainide (F) bound to rat Na_v_1.5 (yellow; PDB code: 6UZ0) and quinidine (Q) bound to human Nav1.5 (orange; PDB code: 6LQA).

Next, we turned to inhibition by flecainide, a slightly larger and more hydrophobic AAD (Fig. 4B). As expected (34), this Class Ic drug showed generally higher apparent affinity than quinidine (Table 2). However, and in contrast to quinidine, we did not observe a phosphorylation-induced change in apparent affinity on the WT background and this was also true for the Q1476R and the ΔK1500 mutations (Fig. 4 B,D and Table 2). Finally, we tested ranolazine, another clinically used sodium channel inhibitor. Like quinidine, ranolazine displayed a reduction in apparent drug affinity on the WT background in response to phosphorylation. This reduction was also observed on the ΔK1500 mutant, but not the Q1476R background (Fig. S5). The above suggests that inhibition of WT Na_v_1.5 by both quinidine and ranolazine is sensitive to phosphorylation at Y1495, while flecainide inhibition is not. Additionally, our data show that the Q1476R mutation abolishes the phosphorylation-induced reduction in apparent affinity for all three tested drugs, while for the ΔK1500 mutation this was only the case for quinidine and flecainide.

## Discussion

In this study, we use a combination of protein semi-synthesis and molecular dynamics simulations to study the effects of phosphorylation on Y1495 on the DIII-DIV linker of Na_v_1.5. Based on this approach, we i) outline a detailed molecular mechanism for how phosphorylation of Y1495 modulates with Na_v_1.5 inactivation, ii) demonstrate that the magnitude of the functional effects caused by this modification can vary substantially when we combine it with a patient mutation and iii) show that phosphorylation can affect the pharmacological sensitivity of Na_v_1.5 towards clinically relevant anti-arrhythmic drugs.

### Mechanism for how Y1495 phosphorylation affects inactivation in Na_v_1.5

Although altered fast inactivation (and late currents) alone can be disease-causing (35) (see also Fig. 3 A-C), it has long been acknowledged that Na_v_1.5 pathogenicity is often associated with changes in the voltage dependence of SSI. For example, the pronounced right-shifted SSI due to phosphorylation of Y1495 is well established from previous *in vitro* work (18, 33). Yet the precise mechanism of how phosphorylation of Y1495 affects SSI, a process fundamental to Na_v_ function, remained enigmatic. Here, we show that the pronounced (~11 mV) right-shift in the voltage sensitivity of SSI and the significantly accelerated recovery from inactivation due to phosphorylation of Y1495 is caused by a destabilization of the interactions between the IFM particle and its receptor site, which results in a decrease of binding energy of the IFM particle to its binding site. Our simulations illustrate that phosphorylation causes the IFM particle to be displaced from its receptor site, moving it towards DIII-S6 to accommodate the phosphate group of Y1495 in the binding pocket of IFM. In light of the high degree of sequence conservation in the DIII-DIV linker across mammalian Na_v_ channels (36) we speculate that the mechanism by which Tyr phosphorylation in the DIII-DIV linker affects channel inactivation is likely conserved beyond Na_v_1.5. It is important to note that Na_v_1.5 SSI likely involves side chains in the C-terminal domain (37–40), possibly including an interaction between the DIII-DIV linker and the channel C-terminus (41). However, our present work is unable to directly address the potential role of phosphorylation in this process.

### Mechanistic insight into Na_v_1.5 disease-causing mutations

The LQT3 mutant Q1476R had previously been shown to leave the voltage dependence of activation unaffected, but to cause a 6.5 mV right-shift in SSI, along with a prominent late current (26). Obtained in HEK293 cells, these findings are consistent with the notion that Na_v_1.5 LQT syndrome mutants typically result in a gain-of-function phenotype (42). We were therefore surprised to find that in the complete absence of phosphorylation at Y1495 (i.e. on the NM background), the Q1476R mutation itself did not change the voltage dependence of SSI, alter the time course of inactivation or elicit a late current (Fig. 2). By contrast, introducing the Q1476R mutation on the phY background caused a dramatic (20 mV) right-shift in SSI, together with a substantial late current and accelerated recovery from inactivation. The more pronounced SSI shift compared to the one observed in HEK293 cells is not unexpected because in our experiments 100% of the Q1476R mutant channels are phosphorylated, whereas native phosphorylation levels under baseline conditions are typically lower (19, 20), which would result in a less pronounced right-shift in SSI.

At the molecular level in our MD simulations, Q1476 appears to stabilize the IFM particle in the WT phY system. The Q1476R mutation, however, disrupts this stabilization and causes significant conformational changes leading to weakened IFM particle binding. The functionally observed right-shift in SSI gives rise to a significant window current around −40 mV (43). Together, our data strongly suggest that the pathogenicity of the LQT3 mutant Q1476R does not arise from the Gln-to-Arg exchange *per se*, but rather the combined effect of mutation and phosphorylation.

To validate the above findings and to ascertain that the observed phosphorylation-induced hyper-shift in SSI was not a nonspecific effect of mutations on the DIII-DIV linker, we also investigated the LQT3 / BS mutant ΔK1500. This mutation has previously been shown to cause a minor right-shift in the voltage dependence of activation, along with a small left-shift in SSI, slower inactivation and a late current (25), similar to what had been observed for e.g. the classic ΔKPQ deletion mutation (44). Our findings show that the ΔK1500 mutation indeed causes a slower time course of inactivation, along with late current on both the NM and phY backgrounds, although the voltage dependence of activation and inactivation were similar to those observed on the WT background. Most importantly, however, we show that the phosphorylation-induced right-shift in SSI is comparable to that of WT (ΔSSI ~12 mV for ΔK1500 vs ~11 mV for WT). This is consistent with our observation that the interaction pattern of the IFM motif with its binding site is no further disrupted than in WT phY, thus underlining that the phosphorylation-induced hyper-shift in SSI is specific to the Q1476R mutation.

### Na_v_1.5 pharmacology can be affected by phosphorylation and patient mutations

Disruptions to inactivation through mutations or other manipulations have previously been shown to weaken binding of local anesthetics to Na_v_ channels (45, 46). This is in line with our finding that quinidine has lower apparent affinity on the WT phY background than on the WT NM background, as the former displays a 11 mV right-shift in SSI. However, on channels with the Q1476R or ΔK1500 mutations, phosphorylation of Y1495 did not lower the apparent affinity towards quinidine, despite phosphorylation causing similar (ΔK1500) or even greater (Q1476R) right-shifts in SSI on these channels compared to WT. This suggests that indirect effects on apparent affinity of Class Ia drugs caused by patient mutations can override those caused by altered inactivation.

Interestingly, we find that on the WT background, inhibition by quinidine and ranolazine is sensitive to phosphorylation, while that by flecainide is not. At least for quinidine and flecainide, where structural data is available, this difference might be explained by their divergent binding positions and poses (11, 13): while flecainide interacts primarily with S6 of DII and DIII, quinidine interacts most closely with S6 of DIV (Fig. 4E). Because the phosphorylated side chain of Y1495 points towards the latter, we speculate that phosphorylation induces an unfavorable conformational change that is most prominent around the quinidine binding site. By contrast, the more drastic conformational consequences of the Q1476R mutation appear to be extensive enough to eliminate any phosphorylation-induced alterations in apparent drug sensitivity.

Although the differences in apparent drug affinity observed by us are comparatively small, our data support the notion that both phosphorylation and disease-causing mutations can impact Na_v_1.5 pharmacology. While disease-causing mutations have been shown previously to affect Na_v_ channel pharmacology (26, 34, 47–49), there is less evidence for phosphorylation-mediated alterations (50). In the future, this may have implications for treatment regimens of patients with known or suspected disease-causing Na_v_1.5 mutations, as some AADs are known to display potentially dangerous pro-arrhythmic properties in some patient subpopulations (i.e. flecainide and encainide) (51–53).

### Potential physiological and clinical relevance of phosphorylation-induced effects

To date, most confirmed Na_v_1.5 phosphorylation sites are situated within the DI-DII linker. Although multiple lines of evidence suggest that all intracellular linkers can be phosphorylated (18, 54, 55), mass spectrometry-based work on cardiac and neuronal Na_v_ channels from native tissues have yet to find evidence for phosphorylation in the DIII-DIV linker (20, 56, 57). This is most likely due to the close association of the DIII-DIV linker with the TM helices (11) and its highly basic sequence content (12/53 aa are basic), both of which reduce MS sequence coverage. By contrast, *in vitro* studies have provided direct evidence for phosphorylation of Y1495 by Fyn kinase (33, 58), as well as phosphorylation of multiple other Tyr side chains in Na_v_1.5 (59). Additionally, tyrosine kinase inhibitors inhibit sodium currents in rabbit cardiac myocytes by left-shifting SSI (60), in line with findings by us and others (23, 33).

The fact that phosphorylation of Y1495 can impact Na_v_1.5 channel availability (Fig. 1), and that this can be exaggerated by patient mutations (Fig. 2), has potential pathophysiological relevance. This is because Tyr kinases have been shown to be more active under pathological conditions, such as ischemia, reperfusion injury or cardiac remodeling (61–63). This aspect is further underlined by the observation that β-adrenergic stimulation causes cardiac ion channels, including Na_v_1.5, to undergo increased phosphorylation (20). Here, we demonstrate that the Q1476R only causes a severe alteration in channel function when phosphorylated. Together, this may help explain why the clinical phenotype of patients with Na_v_1.5 mutations can vary over time or why individuals with the same mutation are affected to different degrees: in the case of Q1476R, the channel is likely to behave WT-like under basal conditions, but increased phosphorylation levels as a result of metabolic alterations and/or disease states would result in a potentially life-threatening shift in SSI.

Therefore, our data suggest that kinase inhibitors could be a treatment option superior to that of traditional AADs for mutations such as Na_v_1.5 Q1476R (54). Additionally, Na_v_1.5 regulation by calmodulin is at least partly mediated through the DIII-DIV linker (64, 65), thus raising the possibility that the interplay of phosphorylation at Y1495 with patient mutations might extend to Ca^2+^ regulation of Na_v_1.5. Lastly, our work emphasizes that it may be beneficial to tailor future AAD treatment regimens towards the specific genetic background of the patient (49).

### Limitations of our study

To ensure efficient delivery of the synthetic peptides, we used microinjection into *Xenopus laevis* oocytes for our tPTS approach. While this affords unique ‘on-and-off’-type control over the extent of phosphorylation at a site of interest (0 *vs* 100%), we cannot assess the extent to which other PTMs might be present in different parts of the protein, or if these would differ in a more relevant mammalian expression system. This is important because the functional impact of Na_v_1.5 mutations can be cell-type specific (66–68). Also, Tyr phosphorylation is typically less abundant than that of Ser/Thr (69), so our findings are likely to overestimate the extent to which phosphorylation of Y1495 would affect Na_v_1.5 function *in vivo*. Similarly, there are minor differences in pKa between the phospho<u>n</u>ylation used in the functional experiments and the phospho<u>r</u>ylation used for the *in silico* work (7.5-8 vs 6.5 (31, 70)). As template for our computational work, we used a Nav1.5 structure that was resolved in a presumably inactivated state. This functional assignment is primarily based on the fact that the IFM particle is docked to a site between the DIII S4-S5 linker and DIV S6. However, we note that the construct used for Na_v_1.5 structure determination was heavily engineered, resulting in severe gating shifts and thus prompting us to carefully question this functional assignment. Despite these caveats, our computational data are in remarkable agreement with the functional characterization of the phosphorylation and the patient mutations, leading us to confidently propose a mechanistic model for the effects caused by these modifications.

## Conclusion

Our study highlights the power of semi-synthetic approaches to decipher complex biophysical interactions. Specifically, the data call for caution when interpreting the apparent functional effects of potentially pathogenic mutations because, at least in some cases, the functional consequences of PTMs may outweigh those caused by mutations. This adds another layer of complexity to the investigation and interpretation of Na_v_1.5 patient mutations, because PTM levels are subject to variation. Given the enormous (>500) number of disease-associated mutations in Na_v_1.5 alone (1), our findings are likely to be relevant beyond the confines of this study. Our work further motivates the investigation of thus far unexplored crosstalk between PTMs and pathological mutations in other proteins, particularly because PTM-mediated effects may not be confined to spatially close regions of the protein. Finally, the results of our study provide a starting point for understanding how Na_v_1.5 mutations can impact the patient-specific pharmacological sensitivity that is observed clinically.

## Methods

### Molecular biology and chemicals

Gene constructs were generated and used as described in (23), see also below for sequence details. Standard site-directed mutagenesis was performed using PCR, deletions were created using the Q5^®^ site-directed mutagenesis kit (Thermo Fisher Scientific). The cDNA was linearized and transcribed into complementary RNA (cRNA) for oocyte microinjection using the Ambion mMESSAGE mMACHINE T7 Transcription Kit (Thermo Fisher Scientific). All chemicals were purchased from Sigma Aldrich unless specified otherwise. Antiarrhythmic drugs were purchased from Sigma-Aldrich (Quinidine sulfate salt dihydrate, Cat #: Q0875; Mexiletine hydrochloride, Cat #: M2727; Flecainide acetate salt, Cat #: F6777; Ranolazine dihydrochloride, Cat #: R6152) and stored according to product specification. Substances were dissolved in ND96 solution (see below), diluted to the specified concentrations and the pH value was adjusted to 7.4. The prepared solutions were then stored at room temperature and used for no longer than 20 days.

### Peptide synthesis

Peptides for Na_v_1.5 splicing were synthesized by solid-phase peptide synthesis as previously described in detail (23). Briefly, P^SYN^ variants were synthesized by ligating three shorter fragments: the N-terminal intein half (Int^C^-A), the sequence derived from the Na_v_1.5 ion channel and the C-terminal intein half (Int^N^-B), with the Na_v_1.5 ion channel sequence being the only variable one. Regarding the ligation strategy adopted, the three fragments were ligated in a ‘one-pot’ fashion and in a C-to-N direction, exploiting the Thz masking group as previously established (71). Note that the WT NM and the phY WT peptides also contained the K1479R mutation (23). Unless otherwise stated, the amino acids used for solid phase peptide synthesis were: Fmoc-Ala-OH; Fmoc-Cys(Trt)-OH; Fmoc-Phe-OH; Fmoc-Gly-OH; Fmoc-Ile-OH; Fmoc-Lys(Boc)-OH; Fmoc-Leu-OH; Fmoc-Pro-OH; Fmoc-His(Trt)-OH; Fmoc-Asn(Trt)-OH; Fmoc-Gln(Trt)-OH; Fmoc-Arg(Pbf)-OH; Fmoc-Ser(*t*Bu)-OH; Fmoc-Thr(*t*Bu)-OH; Fmoc-Tyr(*t*Bu)-OH; Fmoc-Asp(*t*Bu)-OH; Fmoc-Glu(*t*Bu)-OH; Fmoc-Met-OH; Fmoc-Val-OH; Boc-Thz-OH.

Automated peptide synthesis was carried out on a Biotage Syro Wave™ peptide synthesizer using standard Fmoc/tBu SPPS chemistry as previously reported (23). Fmoc deprotection was performed by treatment with piperidine–DMF–formic acid (25:75:0.95, v/v/v), 3 min + 12 min. Coupling reactions were performed as double couplings using Fmoc-Xaa-OH (6.0 equiv to the resin loading), HCTU (6.0 equiv) and *N,N*-diisopropylethylamine (*i*-Pr_2_NEt, 12 equiv) for 40 min for each coupling. All reagents and amino acids used during SPPS were dissolved in *N,N*-dimethylformamide but *i*-Pr_2_NEt, which was dissolved in *N*-methyl-2-pyrrolidinone (NMP). Coupling reactions for non-standard Fmoc-protected amino acids were performed as outlined for each peptide (see Supporting Information). Thioesterification of peptides was performed as described (23, 72) and upon cleavage of protected peptides from resin with 1,1,1,3,3,3-hexafluoro-2-propanol (HFIP) – dichloromethane (20:80, v/v). General deprotection of the peptides was performed with a mixture of trifluoroacetic acid–2,2’-(ethylenedioxy)diethanethiol–triisopropylsilane (TFA– DODT–TIPS, 94:3.3:2.7, v/v/v) for 60–90 min. Upon full deprotection (monitored by MALDI-TOF), the reaction mixture was concentrated under a stream of nitrogen and the crude peptide was precipitated by addition of cold diethyl ether. The solid was spun down and subsequently washed with cold diethyl ether (2x). In case of partial oxidation of methionine residues, this could be reversed by incubating the peptide in a mixture of DODT (0.2 M) and bromotrimethylsilane (TMSBr, 0.1 M) in TFA for 20 min (23, 73). The peptide was then precipitated from diethyl ether before proceeding with preparative HPLC purification.

### Expression in Xenopus laevis oocytes

Stage V-VI oocytes from *Xenopus laevis* oocytes (prepared as described previously (23)) were injected with cRNAs and incubated at 18 °C in OR-3 solution (50 % Leibovitz’s medium, 1 mM L-Glutamine, 250 mg/L Gentamycin, 15 mM HEPES, pH 7.6) for up to 3 days. The lyophilized synthetic peptides were dissolved in Milli-Q H_2_O to a concentration of 500-750 μM and 9-14 nL of solubilized peptide was injected into oocytes pre-injected with cRNA using a *Nanoliter 2010* micromanipulator (World Precision Instruments). Typically, synthetic peptides were injected about 24 hrs after cRNAs injection. Recordings were conducted 12-20 hrs after injection of synthetic peptides.

Note that experimental uncertainties associated with the injection of small amounts of synthetic peptides do not allow us to draw conclusions on expression levels, e.g. to directly compare current sizes between WT and mutant channel variants.

### Two-electrode voltage clamp (TEVC) recordings

Voltage-dependent currents were recorded with two-electrode voltage clamp using an OC-725C voltage clamp amplifier (Warner Instruments). Oocytes were constantly perfused with ND96 solution (in mM: 96 NaCl, 2 KCl, 1 MgCl_2_, 1.8 CaCl_2_ /BaCl_2_, 5 HEPES, pH 7.4) during recordings. For pharmacological testing, solutions containing different antiarrhythmic drugs in ND96 were used. Glass microelectrodes with resistances between 0.1 and 1 MΩ were backfilled with 3 M KCl. To determine values for half-maximal activation, oocytes were held at −100 mV and sodium currents were elicited by voltage steps from −80 mV to +40 mV (in 5 mV increments). Steady-state inactivation curves were determined by applying a 500 ms prepulse from −100 mV to −20 mV (in +5 mV increments), followed by a 25 ms test pulse to −20 mV.

### Molecular dynamics simulations

#### Model building

First, the Na_v_1.5 channel (PDB ID 6UZ3) (11) was embedded in a homogenous lipid bilayer consisting of 400 POPC using the CHARMM-GUI Membrane builder (74). Four different systems were prepared: 1) wild type Nav1.5, 2) Na_v_1.5 with Y1495 phosphorylated, 3) Na_v_1.5 with Y1495 phosphorylated and Q1476R mutation and 4) Na_v_1.5 with Y1495 phosphorylated and ΔK1500 mutation. Phosphorylation and mutation were performed using the CHARMM-GUI Membrane builder. The system was hydrated by adding two ~25 Å layers of water to both sides of the membrane. Lastly, the system was ionized with 150 mM NaCl.

#### Simulations

The Charmm36 force field was used to describe interactions between protein (75), lipid (76) and ions, the TIP3P model was used to describe the water particles (77). The systems were minimized for 5000 steps using steepest descent and equilibrated with constant number of particles, pressure and temperature (NPT) for at least 36 ns for all the four systems, during which the position restraints were gradually released according to the default CHARMM-GUI protocol (78). During equilibration, a time step of 2 fs was used, pressure was maintained at 1 bar through Berendsen pressure coupling, temperature was maintained at 300 K through Berendsen temperature coupling (79) with the protein, membrane and solvent coupled and the LINCs algorithm (80) was used to constrain the bonds involving hydrogen atoms. For long range interactions, periodic boundary conditions and particle mesh Ewald (PME) were used (81). For short range interactions, a cut-off of 12 Å was used. Finally, unrestrained production simulations were run for 200 ns for each of the systems, using Parinello-Rahman pressure coupling (82) and Nose-Hoover temperature coupling (83). Simulations were performed using GROMACS 2019.3 (84, 85).

#### Umbrella Sampling

To calculate the free energy profile of binding of IFM to its binding site, Umbrella sampling was employed, using the distance between the center of mass (COM) of F1486 and the COM of N1659 as a reaction coordinate. Each Umbrella Sampling window was 1 Å wide and the position of the IFM particle was restrained by applying a harmonic potential on the reaction coordinate. A brief 100 ps long NPT equilibration was conducted with 1 atm pressure maintained by Berendsen pressure coupling, and 300K, controlled by Berendsen thermostat. This was followed by 10 ns of production simulation in each window. The N1659 residue (located in the IFM particle binding site) was restrained throughout the simulation with a restraining potential force constant of 1000 kJ·mol^−1^ ·nm^−2^. The weighted histogram analysis method (WHAM) (86) in gromacs (gmx wham) was used to combine data from all the umbrella sampling windows to compute the potential of mean force (PMF).

#### Contacts

Contacts are defined when the distance between the Cβ atoms of pairs of residues are below 6.7 Å. Those were calculated using MD-TASK (87).

## Acknowledgements

We acknowledge the Lundbeck Foundation (R139-2012-12390), the Independent Research Fund Denmark (7025-00097A and 9039-00335B) (all to SAP), SciLifeLab and the Swedish Research Council to LD (VR 2018-04905) for funding. The MD simulations were performed on resources provided by the Swedish National Infrastructure for Computing (SNIC) at PDC Centre for High Performance Computing (PDC-HPC). We thank Prof Christian A Olsen for support with the peptide chemistry and would like to thank members of the Pless lab for helpful comments on the manuscript.

## Author contributions

I.G., H.H., K.C., K.K.K., L.D. and S.A.P. designed the research. I.G., H.H. K.C. and K.K.K. performed the experiments and analyzed the data. All authors interpreted the data. I.G., L.D. and S.A.P. drafted the manuscript, which was finalized with input from all authors.

