## Supplementary information for "Functional crosstalk between phosphorylation and disease-causing mutations in the cardiac sodium channel Na_v_1.5"

- Supplementary Figures 1-5
- Supplementary Text
- Supplementary References

### Supplementary Figures

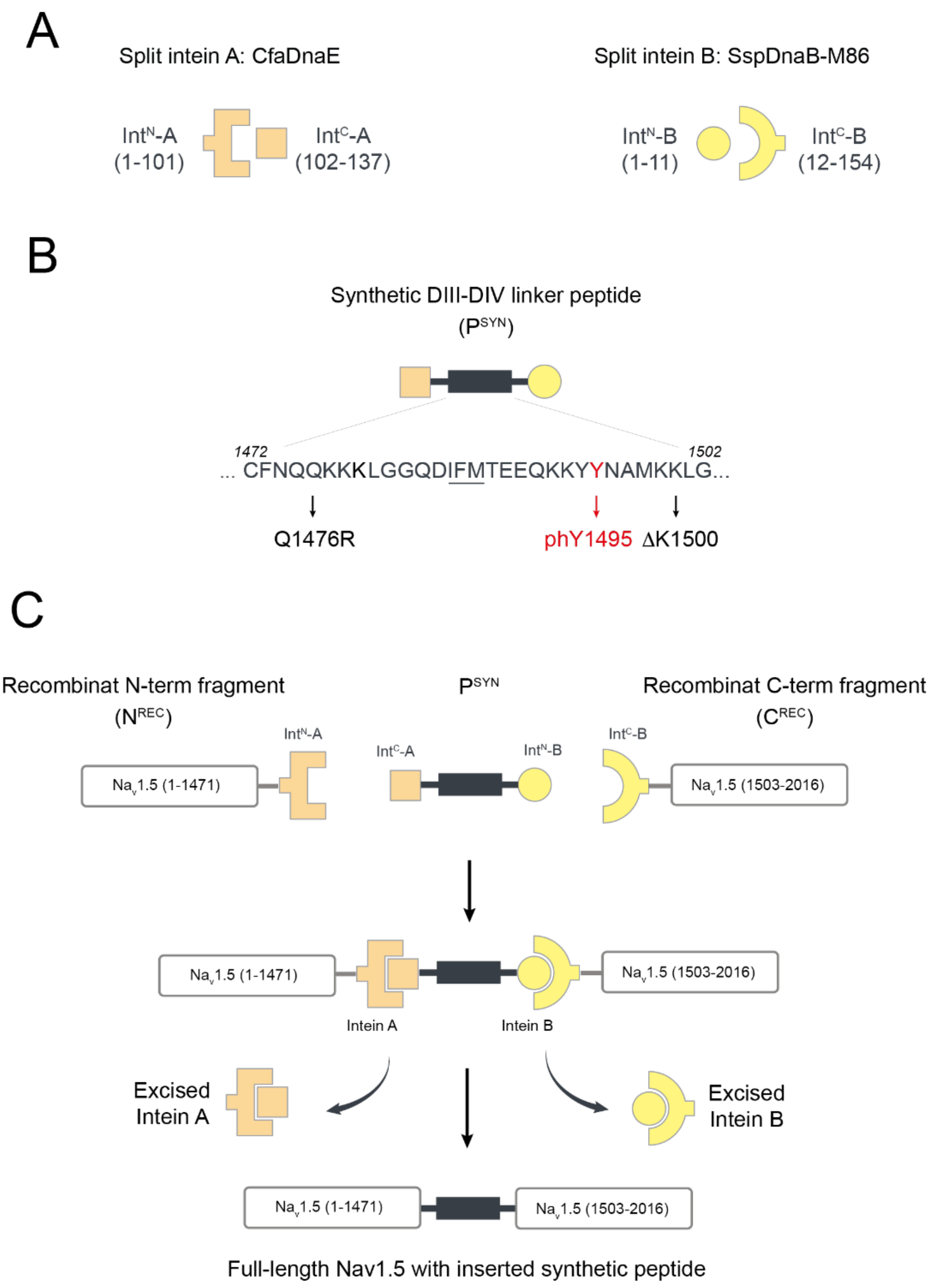


**Fig. S1.** Schematic overview of the tPTS-based approach to generate semi-synthetic $Na_v1.5$ channels. (A) Split intein A (CfaDnaE; orange) and B (SspDnaB M86; yellow), with N- and C-terminal split intein parts shown separately. Numbers in brackets indicate amino acid numbers of split inteins. (B) Sequence of the synthetic peptide ($P^{SYN}$) corresponding to amino acids 1472 to 1502 of $Na_v1.5$. Mutated (Q1476R and ΔK1500) or modified (Y1495) sites are highlighted; the IFM motif sequence is underlined. (C) Summary of the approach, with N- and C-terminal $Na_v1.5$ fragments ($N^{REC}$ and $C^{REC}$) being covalently spliced on both ends of $P^{SYN}$. Numbers in brackets indicate amino acid numbers in $Na_v1.5$.

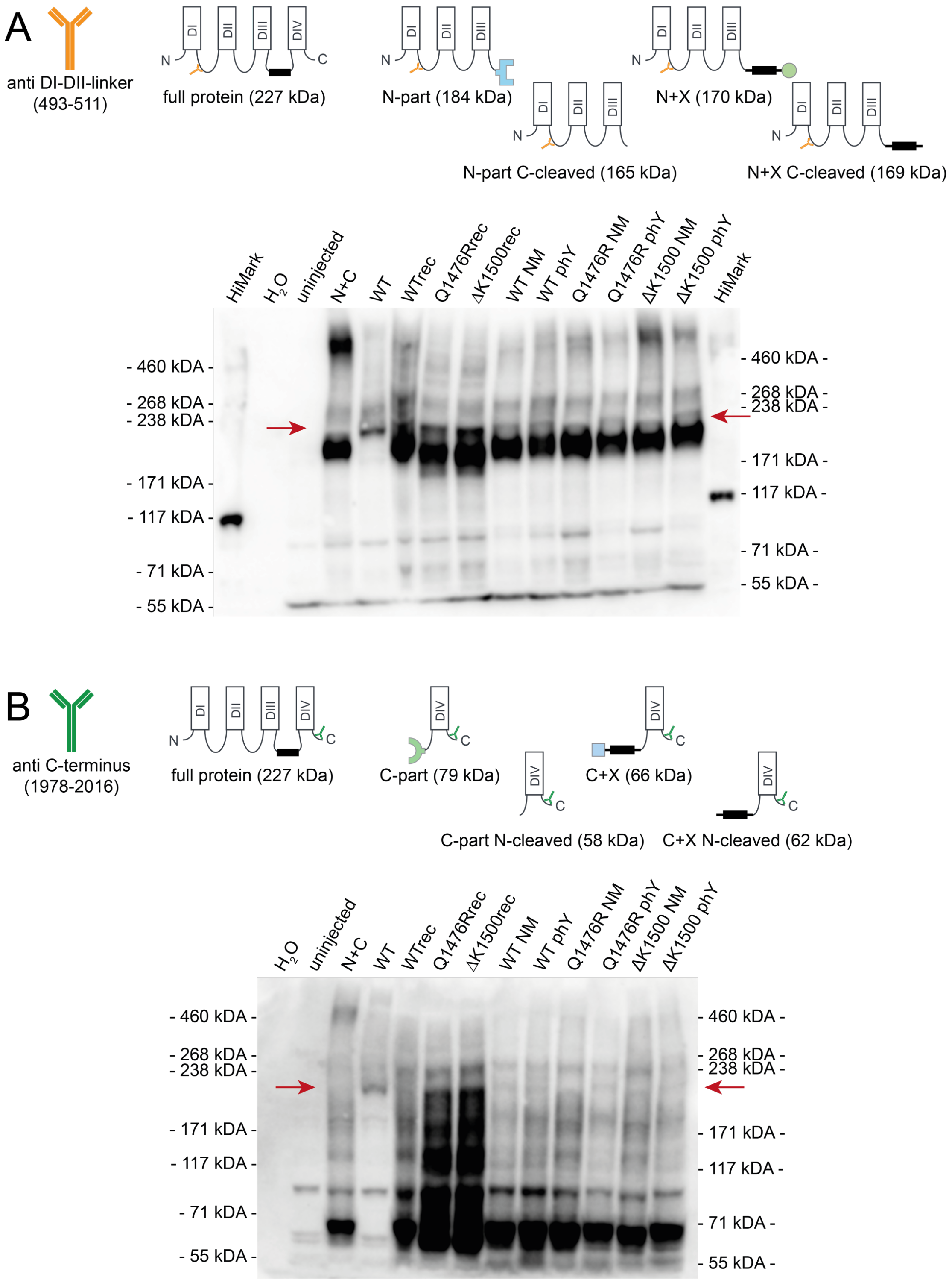


**Fig. S2.** Western blots of full-length Nav1.5 protein and semi-synthetic channel constructs using antibodies directed against Nav1.5 DI-DII-linker (A) or Nav1.5 C-terminus (B). Top: The respective antibody is schematically shown on the left with its respective epitope (hNav1.5 amino acid numbering) denoted in brackets below it. The detectable constructs (fully spliced protein, splicing educts, splicing side products) are shown in basic topology and their mass is indicated in brackets. Bottom: Western blots of whole-cell lysate from *Xenopus laevis* oocytes expressing the respective constructs, using a different epitope. Red arrows mark the area around 230 kDa, corresponding to the mass of full length Nav1.5 protein. Lysate from uninjected oocytes and water instead of cell lysate were used as negative controls. Note that the blots also include two constructs with mutated variants of recombinant peptide X (Q1476Rrec, ΔK1500rec).

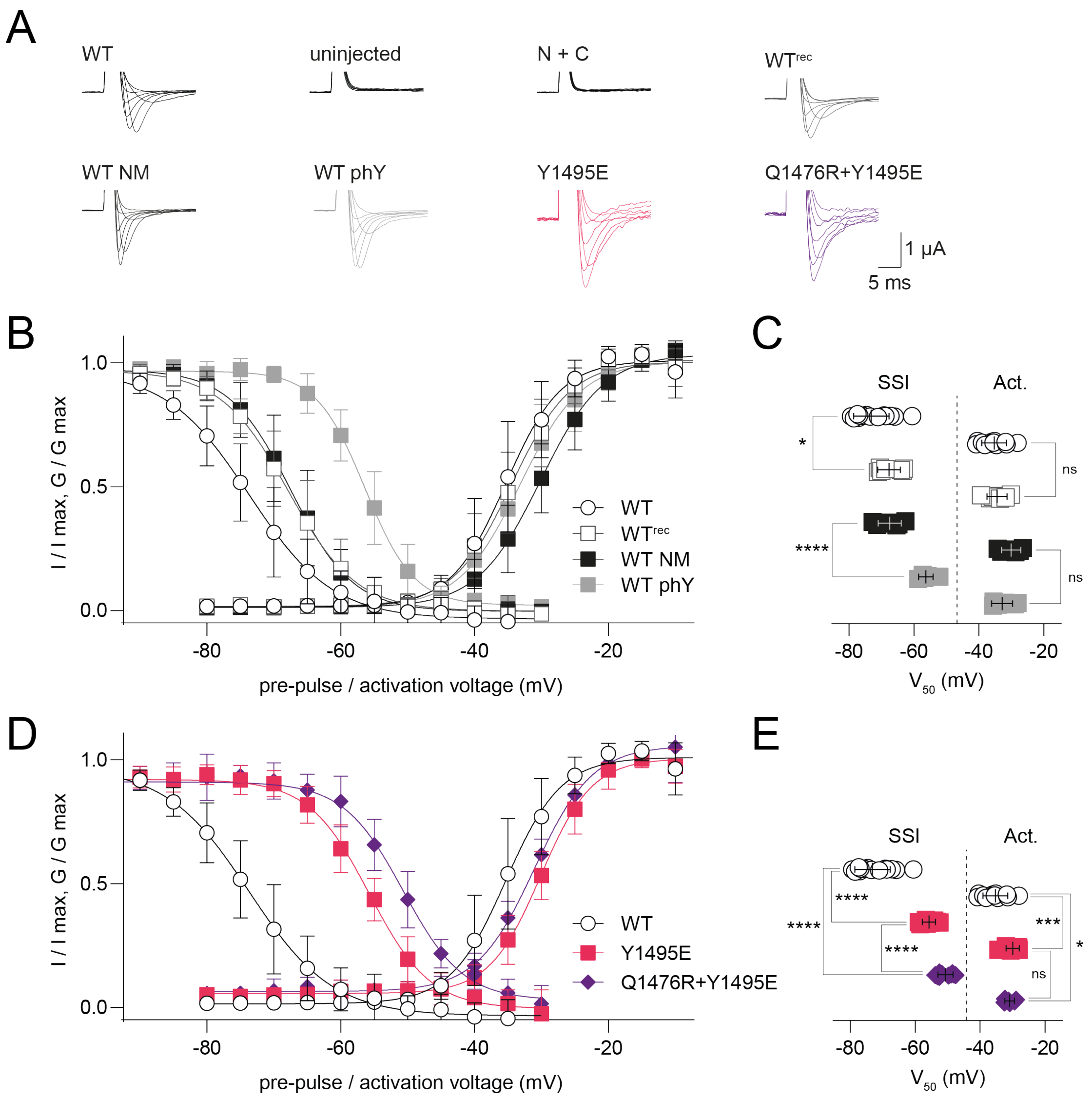


**Fig. S3.** Functional characterization of conventional WT, recombinant reconstituted and semi-synthetic Na$_V$1.5 constructs, as well as conventional single and double mutations. (A) Representative current recordings of *Xenopus laevis* oocytes injected with the indicated constructs. WT = full length WT Na$_V$1.5; WT$^{REC}$ = N$^{REC}$ + C$^{REC}$ + P$^{REC}$; WT NM = N$^{REC}$ + C$^{REC}$ + P$^{SYN}$ NM (i.e. containing the Y1495F and the K1479R mutations); WT phY = N$^{REC}$ + C$^{REC}$ + P$^{SYN}$ phY (i.e. containing a phosphonylated tyrosine in position 1495, in addition to the K1479R mutation); Y1495E, full length Nav1.5 Y1495E single mutant; Q1476R+Y1495E, full length Nav1.5 Q1476R+Y1495E double mutant. Note that uninjected oocytes, or those injected with only N$^{REC}$ and C$^{REC}$ (N + C) did not yield currents. (B) and (D) SSI (left) and activation (right) curves of indicated constructs. (C) and (E) Summary of SSI and activation (Act.) parameters of indicated constructs, see Table 1 for details. Data shown as mean ± standard deviation in (B) - (E); n = 6-16, data was compared using unpaired two-tailed student's t-test, ns (not significant) $p > 0.05$, * $p \leq 0.05$, *** $p \leq 0.001$, **** $p < 0.0001$. Note that the full-length mutant channels were missing the N1472C mutation introduced in the engineered split channels, suggesting that the observed difference in SSI between Q1476R NM and Q1476R phY is likely not an artifact due to the splice-promoting mutation.

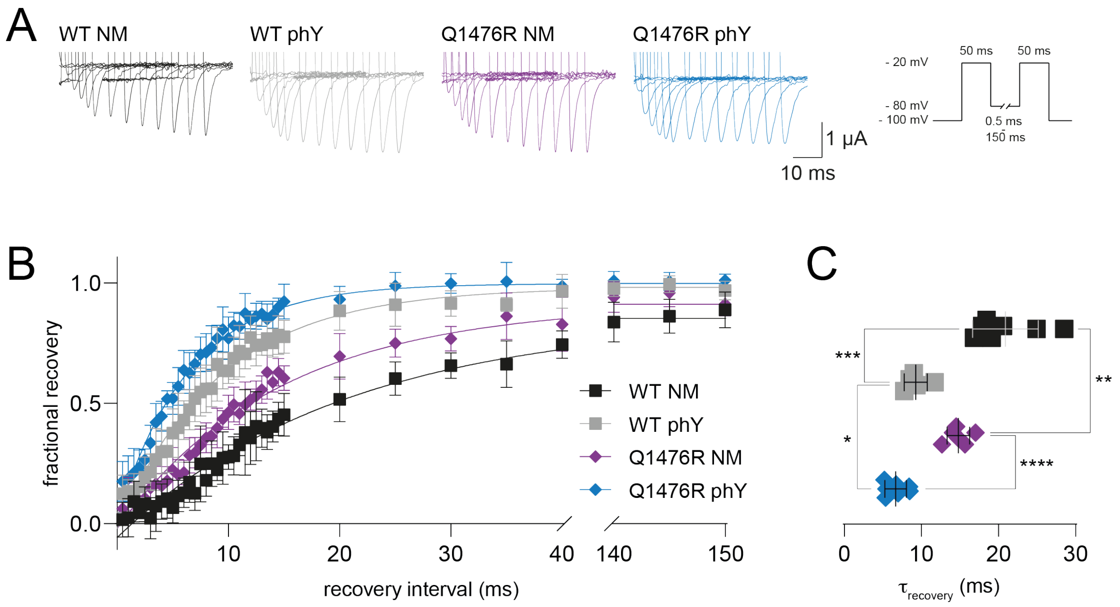


**Fig. S4.** Recovery from inactivation is accelerated by phosphorylation and the Q1476R patient mutation. (A) Representative current recordings of *Xenopus laevis* oocytes expressing the indicated semi-synthetic $Na_v1.5$ constructs. Oocytes were subjected to a voltage protocol consisting of two depolarizing voltage pulses to - 20 mV with a recovery interval at - 80 mV of varying length (0.5 ms to 150 ms) in between (see schematic on the right). Only a selection of traces is shown, and some traces are cropped at the beginning for clarity. (B) Recovery of channels is plotted as a single exponential function over increasing recovery intervals. Fractional recovery is calculated as the peak current elicited by the second depolarizing voltage pulse divided by the peak current elicited by the first depolarizing voltage pulse. Recovery interval is the time between the end of the first and the start of the second depolarizing voltage pulse. (C) Tau values of channel recovery, see Table 1 for details. Data shown as mean ± standard deviation; n = 5-7, data was compared using unpaired two-tailed student's t-test, * p ≤ 0.05, ** p ≤ 0.01, *** p ≤ 0.001, **** p < 0.0001.

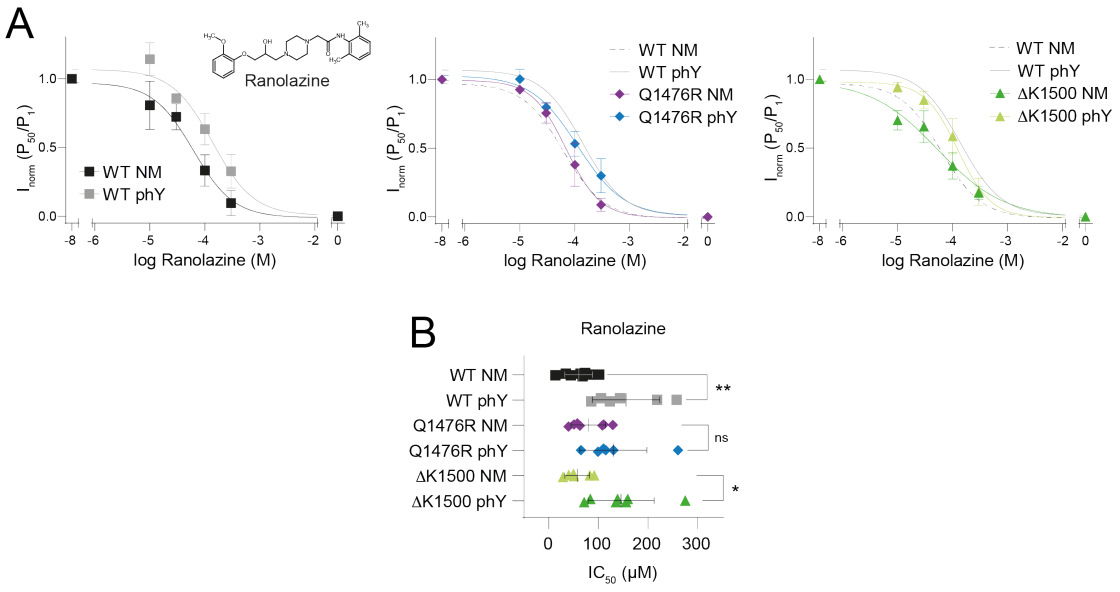


**Fig. S5.** Phosphorylation and disease mutations can affect pharmacological sensitivity of $Na_V1.5$ towards the anti-arrhythmic drug ranolazine. (A) Concentration response curves of WT, Q1476R and ΔK1500 constructs in response to a 20 Hz pulse train stimulation in presence of the antiarrhythmic drug ranolazine (structure shown in left panel). $P_{50}/P_1$ values were normalized to range from 0 to 1. (B) $IC_{50}$ values obtained for ranolazine data shown in (A). The $IC_{50}$ is significantly increased by phosphorylation only in WT and ΔK1500, but not in Q1476R constructs. Data shown as mean ± standard deviation; n = 6-8, data was compared using unpaired two-tailed student's t-test, ns (not significant) $p > 0.05$, * $p \leq 0.05$, ** $p \leq 0.01$.

### Supplementary Text

*Material and Methods*

*Design of plasmid DNA constructs*

Plasmid DNA constructs were designed to encode for the respective protein sequences shown below, see also (1):

<u>Nav1.5 DIII-DIV linker splicing constructs</u>

N-construct

pUNIV - hNav1.5(aa 1-1471) - **CfaDnaE$_{\mathbf{N101}}$** – *HA tag linker* - *<u>ER retention signal</u>*

MANFLLPRGTSSFRRFTRESLAAIEKRMAEKQARGSTTLQESREGLPEEEA
PRPQLDLQASKKLPDLYGNPPQELIGEPLEDLDPFYSTQKTFIVLNKGKTIFR
FSATNALYVLSPFHPIRRAAVKILVHSLFNMLIMCTILTNCVFMAQHDPPPWT
KYVEYTFTAIYTFESLVKILARGFCLHAFTFLRDPWNWLDFSVIIMAYTTEFVD
LGNVSALRTFRVLRALKTISVISGLKTIVGALIQSVKKLADVMVLTVFCLSVFAL
IGLQLFMGNLRHKCVRNFTALNGTNGSVEADGLVWESLDLYLSDPENYLLK
NGTSDVLLCGNSSDAGTCPEGYRCLKAGENPDHGYTSFDSFAWAFLALFR
LMTQDCWERLYQQTLRSAGKIYMIFFMLVIFLGSFYLVNLILAVVAMAYEEQN
QATIAETEEKEKRFQEAMEMLKKEHEALTIRGVDTVSRSSLEMSPLAPVNSH
ERRSKRRKRMSSGTEECGEDRLPKSDSEDGPRAMNHLSLTRGLSRTSMKP
RSSRGSIFTFRRRDLGSEADFADDENSTAGESESHHTSLLVPWPLRRTSAQ
GQPSPGTSAPGHALHGKKNSTVDCNGVVSLLGAGDPEATSPGSHLLRPVM
LEHPPDTTTPSEEPGGPQMLTSQAPCVDGFEEPGARQRALSAVSVLTSALE
ELEESRHKCPPCWNRLAQRYLIWECCPLWMSIKQGVKLVVMDPFTDLTITM
CIVLNTLFMALEHYNMTSEFEEMLQVGNLVFTGIFTAEMTFKIIALDPYYYFQ
QGWNIFDSIIVILSLMELGLSRMSNLSVLRSFRLLRVFKLAKSWPTLNTLIKIIG
NSVGALGNLTLVLAIIVFIFAVVGMQLFGKNYSELRDSDSGLLPRWHMMDFF
HAFLIIFRILCGEWIETMWDCMEVSGQSLCLLVFLLVMVIGNLVVLNLFLALLL
SSFSADNLTAPDEDREMNNLQLALARIQRGLRFVKRTTWDFCCGLLRQRPQ
KPAALAAQGQLPSCIATPYSPPPPETEKVPPTRKETRFEEGEQPGQGTPGD
PEPVCVPIAVAESDTDDQEEDEENSLGTEEESSKQQESQPVSGGPEAPPDS
RTWSQVSATASSEAEASASQADWRQQWKAEPQAPGCGETPEDSCSEGST
ADMTNTAELLEQIPDLGQDVKDPEDCFTEGCVRRCPCCAVDTTQAPGKVW
WRLRKTCYHIVEHSWFETFIIFMILLSSGALAFEDIYLEERKTIKVLLEYADKMF
TYVFVLEMLLKWVAYGFKKYFTNAWCWLDFLIVDVSLVSLVANTLGFAEMG

PIKSLRTLRALRPLRALSRFEGMRVVVNALVGAIPSIMNVLLVCLIFWLIFSIM
GVNLFAGKFGRCINQTEGDLPLNYTIVNNKSQCESLNLTGELYWTKVKVNFD
NVGAGYLALLQVATFKGWMDIMYAAVDSRGYEEQPQWEYNLYMYIYFVIFII
FGSFFTLNLFIGVIID**CLSYDTEILTVEYGFLPIGKIVEERIECTVYTVDKNGFV**
**YTQPIAQWHNRGEQEVFEYCLEDGSIIRATKDHKFMTTDGQMLPIDEIFERG**
**LDLKQVDGLP***YPYDVPDYAYPYDVPDYLLDALTLASSRGPLRKRSVAVAKA*
*KPKFSISPDSLS*PRKKFQ*

P-construct ‘Rec’

pUNIV - **CfaDnaE**$_{C35}$ - hNav1.5(aa 1472-1502, N1472C) - **SspDnaB**$^{M86}_{N11}$

**VKIISRKSLGTQNVYDIGVEKDHNFLLKNGLVASN**CFNQQKKKLGGQDIFMT
EEQKKYYNAMKKLG**CISGDSLISLA**

C-construct

pUNIV – *ER retention signal* - *linker* - **SspDnaB**$^{M86}_{C143}$ - hNav1.5(aa 1503-2016)

*MLLDALTLASSRGPLRKRSVAVAKAKPKFSISPDSLSGSAGSAAGSGEF***STG**
**KRVPIKDLLGEKDFEIWAINEQTMKLESAKVSRVFCTGKKLVYTLKTRLGR**
**TIKATANHRFLTIDGWKRLDELSLKEHIALPRKLESSSLQLAPEIEKLPQSDI**
**YWDPIVSITETGVEEVFDLTVPGLRNFVANDIIVHN**SKKPQKPIPRPLNKYQG
FIFDIVTKQAFDVTIMFLICLNMVTMMVETDDQSPEKINILAKINLLFVAIFTGEC
IVKLAALRHYYFTNSWNIFDFVVVILSIVGTVLSDIIQKYFFSPTLFRVIRLARIG
RILRLIRGAKGIRTLLFALMMSLPALFNIGLLLFLVMFIYSIFGMANFAYVKWEA
GIDDMFNFQTFANSMLCLFQITTSAGWDGLLSPILNTGPPYCDPTLPNSNGS
RGDCGSPAVGILFFTTYIIISFLIVVNMYIAIILENFSVATEESTEPLSEDDFDMF
YEIWEKFDPEATQFIEYSVLSDFADALSEPLRIAKPNQISLINMDLPMVSGDRI
HCMDILFAFTKRVLGESGEMDALKIQMEEKFMAANPSKISYEPITTTLRRKHE
EVSAMVIQRAFRRHLLQRSLKHASFLFRQQAGSGLSEEDAPEREGLIAYVM
SENFSRPLGPPSSSSISSTSFPPSYDSVTRATSDNLQVRGSDYSHSEDLADF
PPSPDRDRESIV*

*Characterization of peptides*

Detailed characterization of Na$_v$1.5 WT NM and Na$_v$1.5 WT phY1495 peptides and the corresponding ligated P$^{SYN}$ variants has been published previously (1). Low resolution mass spectra were recorded on a MALDI-TOF Bruker Microflex LT/SH system, and samples were prepared using SA (sinapic acid) matrix dissolved in water–MeCN–TFA (50:50:0.1, v/v/v). The calculated mass reported is the most intense peak (100% relative intensity), predicted with mMass software.

High resolution mass spectra (HR-MS) were recorded on a SOLARIX ESI MALDI from Bruker Daltronik. Samples were dissolved in MeCN–water–FA (50:50:0.1, v/v/v) and were analyzed by ESI. The calculated mass reported is the most intense peak predicted with mMass software (100% relative intensity) in the isotopes pattern, which is compared to the most intense peak experimentally found in the isotopes pattern. The second most intense peak in the isotope pattern, predicted and experimentally found, is reported in parenthesis.

Analytical reverse-phase HPLC was performed on an Agilent 1100 LC system equipped with a C8 Phenomenex Kinetex column [250 mm x 4.60 mm, 5 µm, 100 Å] and a diode array UV detector, using a gradient and rising eluent II (0.1% TFA in MeCN) in eluent I (water MeCN–TFA, 95:5:0.1, v/v/v) linearly from 0% to 40% over 40 min, with a flow rate of 1.2 mL/min at 40 °C.

Preparative reversed-phase HPLC was performed on an Agilent 1260 Infinity system equipped with a C8 Phenomenex Luna column [250 mm x 21.2 mm, 5 µm, 100 Å] and a diode array UV detector, using a gradient of eluent I (water–MeCN–TFA, 95:5:0.1, v/v/v) and eluent II (0.1% TFA in MeCN) as specified for each compound, with a flow rate of 20 mL/min.

*$Na_v$1.5_Q1476R_NM*

The peptide was synthesized on pre-loaded Fmoc-Gly Trityl TentaGel resin according to the general peptide synthesis protocol outlined in the Methods section. HPLC purification yielded a white, fluffy solid (7.9 mg as a TFA salt; yield 8% from a 20 µmol scale synthesis).

Prep-HPLC purification conditions: 0–15% eluent II in eluent I (5 min gradient) followed by 15–35% eluent II in eluent I (30 min gradient).

Low resolution MS (MALDI-TOF): calc. $[C_{171}H_{275}N_{46}O_{46}S_4]^+$ $[M + H]^+$: 3837.9 Da; found: 3836.4 Da.

HR-MS: calc. $[M + 5H]^{5+}$: 768.3959 Da (768.5966 Da); found: 768.5971 Da (768.7973 Da)

calc. $[M + 6H]^{6+}$: 640.4978 Da (640.6650 Da); found: 640.6657 Da (640.8325 Da)

calc. $[M + 7H]^{7+}$: 549.1420 Da (549.2853 Da); found: 549. 2858 Da (549.1427 Da)

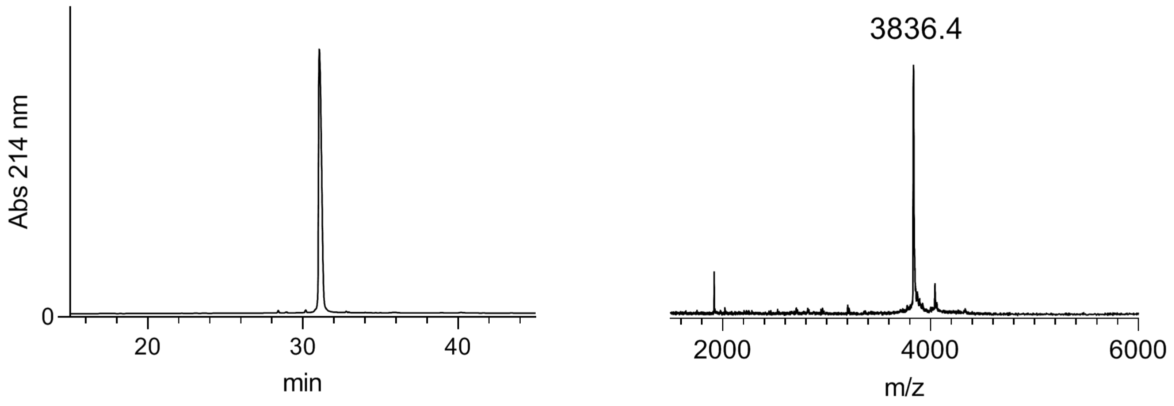


*Na$_v$1.5_Q1476R_phY1495*

The peptide was synthesized on pre-loaded Fmoc-Gly Trityl TentaGel resin according to the general peptide synthesis protocol outlined in the Methods section. Fmoc-Pmp(*t*Bu)$_2$-OH (phosphonylated tyrosine building block) was introduced in the growing peptide with a single coupling (2 h) using 2.5 equiv, HATU (2.5 equiv) and *i*-Pr$_2$NEt (5.0 equiv). HPLC purification yielded a white, fluffy solid (8.0 mg as a TFA salt; yield 8% from a 20 µmol scale synthesis).

Prep-HPLC purification conditions: 0–15% eluent II in eluent I (5 min gradient) followed by 15–35% eluent II in eluent I (30 min gradient).

Low resolution MS (MALDI-TOF): calc. $[C_{172}H_{278}N_{46}O_{49}PS_4]^+$ $[M + H]^+$: 3931.9 Da; found: 3931.0 Da.

HR-MS: calc. $[M + 5H]^{5+}$: 787.1923 Da (787.3930 Da); found: 787.3921 Da (787.1916 Da)

calc. $[M + 6H]^{6+}$: 656.1615 Da (656.3287 Da); found: 656.3281 Da (656.1609 Da)

calc. $[M + 7H]^{7+}$: 562.7113 Da (562.5680 Da); found: 562.7110 Da (562.5677 Da)

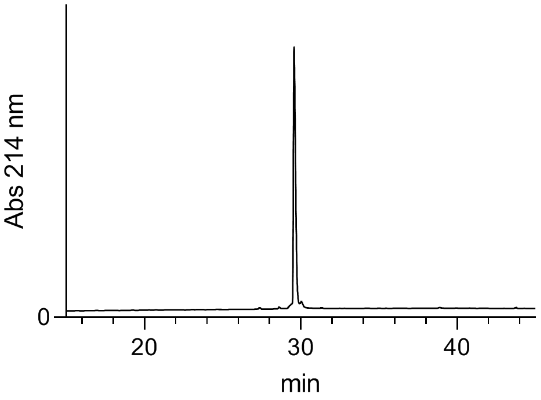


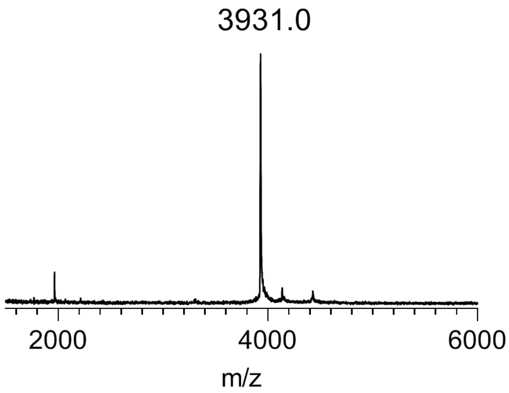


*$Na_v$1.5_ΔK1500_NM*

The peptide was synthesized on pre-loaded Fmoc-Gly Trityl TentaGel resin according to the general peptide synthesis protocol outlined in the Methods section. HPLC purification yielded a white, fluffy solid (6.7 mg as a TFA salt; yield 7% from a 20 µmol scale synthesis).

Prep-HPLC purification conditions: 0–15% eluent II in eluent I (5 min gradient) followed by 15–35% eluent II in eluent I (30 min gradient).

Low resolution MS (MALDI-TOF): calc. $[C_{164}H_{259}N_{42}O_{46}S_4]^+$ $[M + H]^+$: 3681.8 Da; found: 3681.0 Da.

HR-MS: calc. $[M + 5H]^{5+}$: 737.1684 Da (737.3691 Da); found: 737.3683 Da (737.1679 Da)

calc. $[M + 6H]^{6+}$: 614.4749 Da (614.6421 Da); found: 614.6415 Da (614.4745 Da)

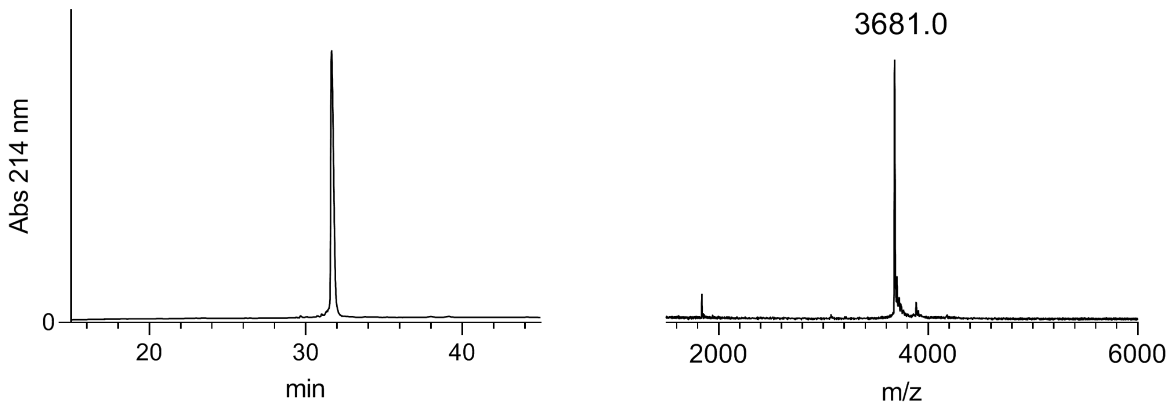


*Na$_V$1.5_ΔK1500_phY1495*

The peptide was synthesized on pre-loaded Fmoc-Gly Trityl TentaGel resin according to the general peptide synthesis protocol outlined in the Methods section. Fmoc-Pmp(*t*Bu)$_2$-OH (phosphonylated tyrosine building block) was introduced in the growing peptide with a single coupling (2 h) using 2.5 equiv, HATU (2.5 equiv) and *i*-Pr$_2$NEt (5.0 equiv). HPLC purification yielded a white, fluffy solid (6.4 mg as a TFA salt; yield 7% from a 20 μmol scale synthesis).
Prep-HPLC purification conditions: 0–15% eluent II in eluent I (5 min gradient) followed by 15–35% eluent II in eluent I (30 min gradient).
Low resolution MS (MALDI-TOF): calc. $[C_{165}H_{262}N_{42}O_{49}PS_4]^+$ $[M + H]^+$: 3775.8 Da; found: 3777.3 Da.
HR-MS: calc. $[M + 5H]^{5+}$: 755.9648 Da (756.1655 Da); found: 756.1655 Da (755.9653 Da)
calc. $[M + 6H]^{6+}$: 630.1386 Da (630.3058 Da); found: 630.3056 Da (630.1388 Da)

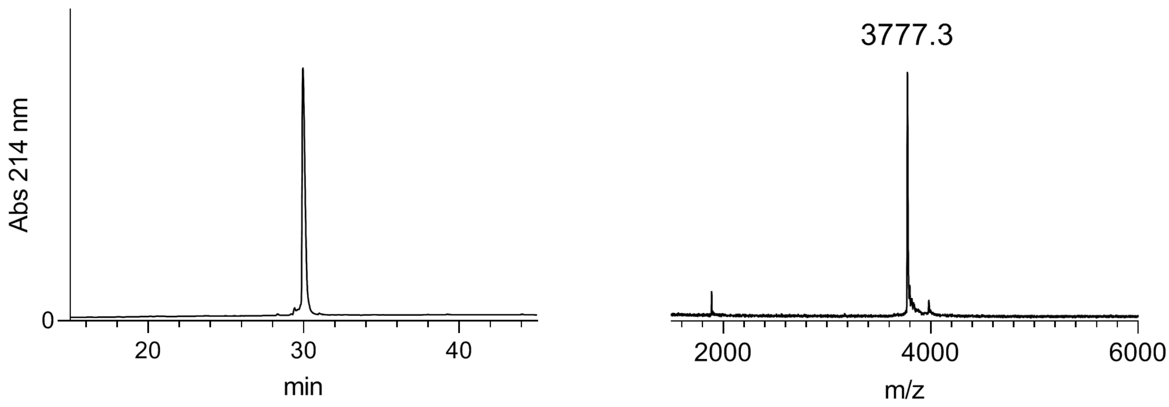


*General procedure for one-pot 'C-to-N directed' ligations*

Reactions were performed in ligation buffer (guanidinium chloride 6M, $Na_2HPO_4$ 100mM, pH 8), which was sparged with nitrogen before use. Tris(2-carboxyethyl)phosphine hydrochloride (TCEP·HCl) was then added (20 mM), followed by addition of 2,2,2-trifluoroethanethiol (TFET) (2% v/v) (2). The pH was adjusted to ~7 with NaOH (5 M) before adding the ion channel peptide fragment (0.57 µmol, 1.0 equiv, 2 mM), followed by the $Int^N$-B peptide fragment (0.57 µmol, 1.0 equiv). The pH was readjusted to ~7 and the reaction mixture incubated at 37 °C. After conversion to the desired ligated product (monitored by HPLC and MALDI-TOF), TCEP·HCl (to reach 40 mM final concentration) and methoxylamine hydrochloride ($MeONH_2$·HCl, to reach 200 mM final concentration) were dissolved in buffer (15 µL) and added to the ligation mixture, which was then incubated at 37 °C. After conversion to the desired unmasked N-terminal cysteine peptide (monitored by HPLC and MALDI-TOF), the pH was readjusted to ~7. TFET (1% v/v) and the $Int^C$-A peptide fragment (0.57 µmol, 1.0 equiv) were then added. The pH was adjusted to ~7 and the reaction was incubated at room temperature. After over night incubation, the ligation mixture was subjected to preparative HPLC purification.

$P^{SYN}_{Na_V1.5\_Q1476R\_NM}$

The full peptide was assembled according to the general procedure 'C-to-N directed ligations' described above. Preparative HPLC purification followed by lyophilization yielded the peptide as a fluffy solid (1.6 mg as a TFA salt; yield 27 %).

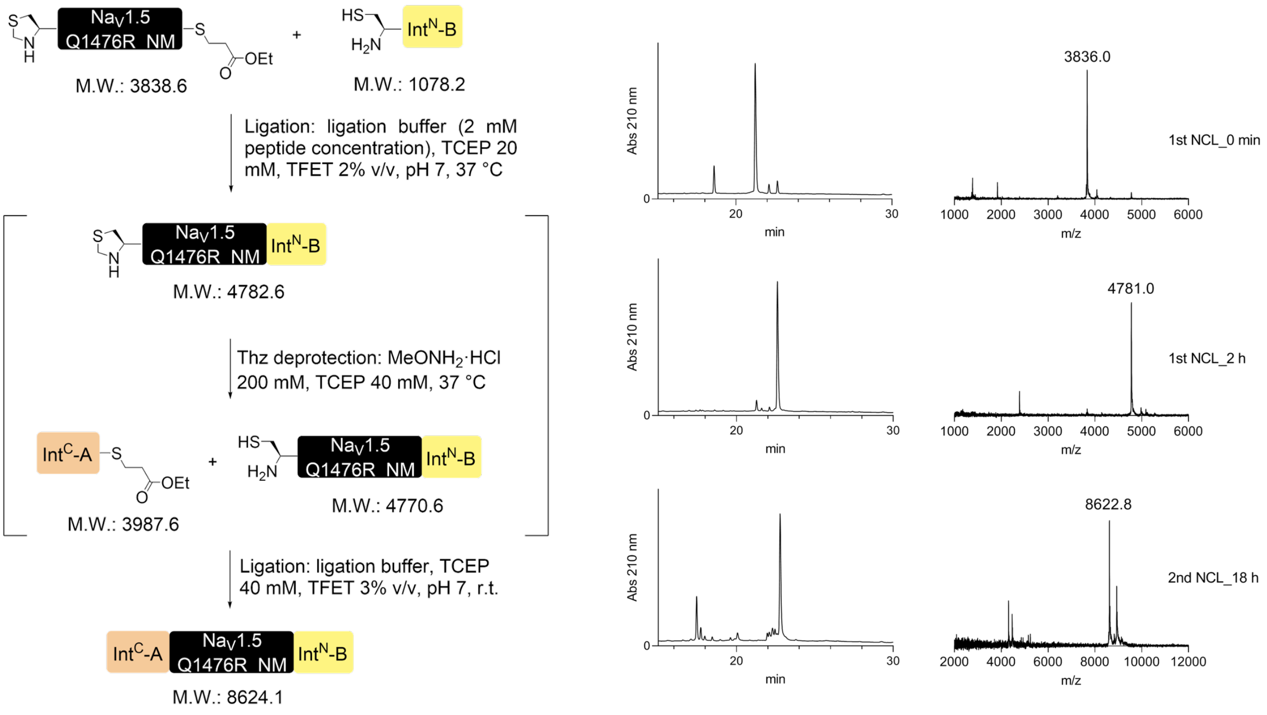


Prep-HPLC purification conditions: 0–10% eluent II in eluent I (5 min gradient) followed by 10–45% eluent II in eluent I (45 min gradient).

Low resolution MS (MALDI-TOF): calc. $[C_{382}H_{627}N_{106}O_{112}S_4]^+$ $[M + H]^+$: 8623.6 Da; found: 8623.6 Da.

HR-MS: calc. $[M + 9H]^{9+}$: 959.0690 Da (959.1804 Da); found: 959.1812 Da (959.0698 Da)

calc. $[M + 10H]^{10+}$: 863.2628 Da (863.3631 Da); found: 863.3639 Da (863.4643 Da)

calc. $[M + 11H]^{11+}$: 784.8759 Da (784.9671 Da); found: 784.9680 Da (784.8770 Da)

calc. $[M + 12H]^{12+}$: 719.5535 Da (719.6371 Da); found: 719.6376 Da (719.5543 Da)

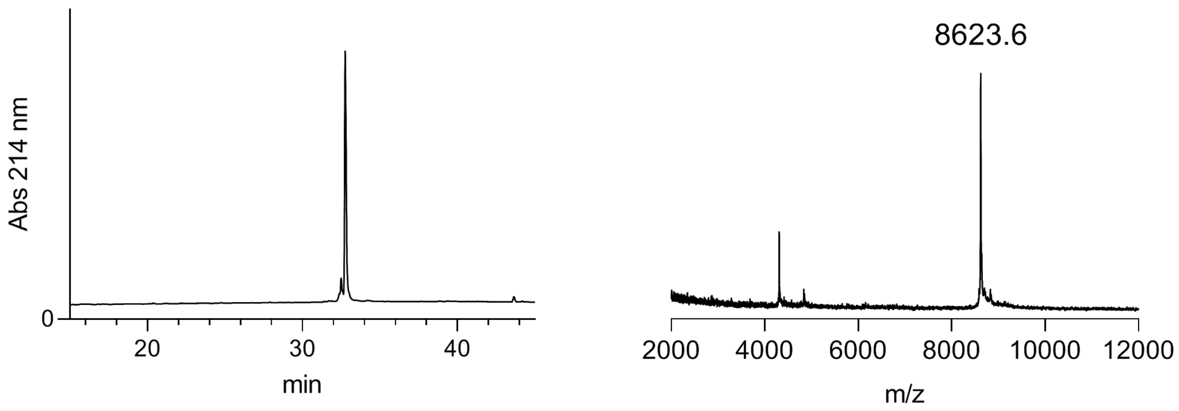


*$P^{SYN}$ $_{Na_V1.5\_Q1476R\_phY1495}$*

The full peptide was assembled according to the general procedure ‘C-to-N directed ligations’ described above, except that the ion channel peptide fragment (Na$_V$1.5_Q1476R_phY1495) was used in slight excess (5.98*10$^{-7}$ mol, 1.05 equiv) in the first step of the ligation. Preparative HPLC purification followed by lyophilization yielded the peptide as a fluffy solid (1.6 mg as a TFA salt; yield 27%).

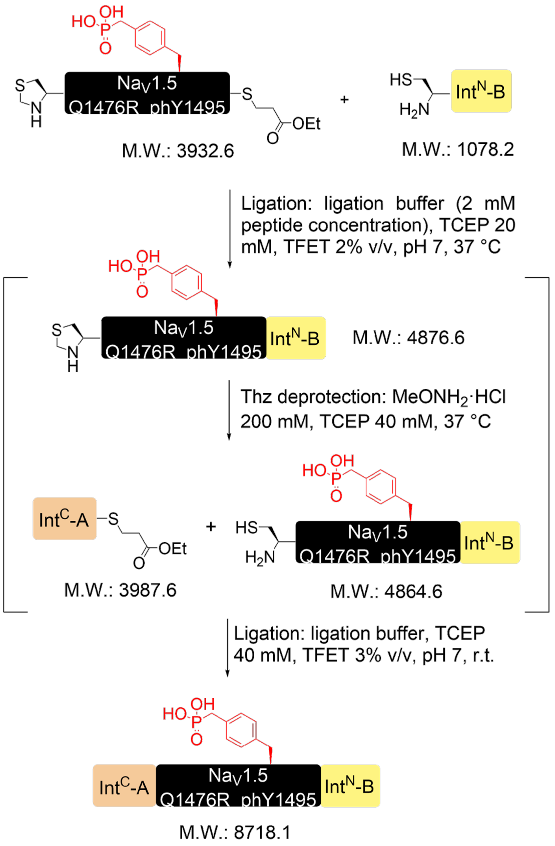


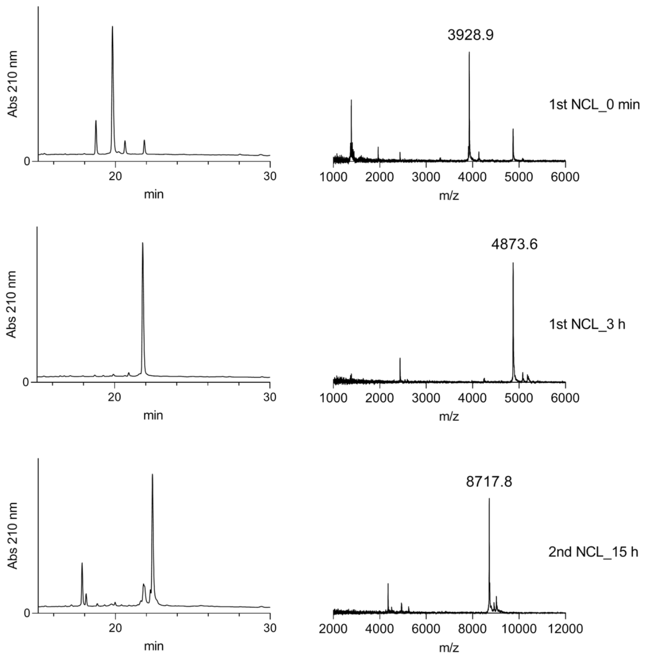


Prep-HPLC purification conditions: 0–10% eluent II in eluent I (5 min gradient) followed by 10–45% eluent II in eluent I (45 min gradient).

Low resolution MS (MALDI-TOF): calc. $[C_{383}H_{630}N_{106}O_{115}PS_4]^+$ $[M + H]^+$: 8717.5 Da; found: 8716.9 Da.

HR-MS: calc. $[M + 9H]^{9+}$: 969.5114 Da (969.6229 Da); found: 969.6226 Da (969.5112 Da)

calc. $[M + 11H]^{11+}$: 793.4197 Da (793.5109 Da); found: 793.5105 Da (793.4196 Da)

calc. $[M + 12H]^{12+}$: 727.3853 Da (727.4689 Da); found: 727.4687 Da (727.3852 Da)

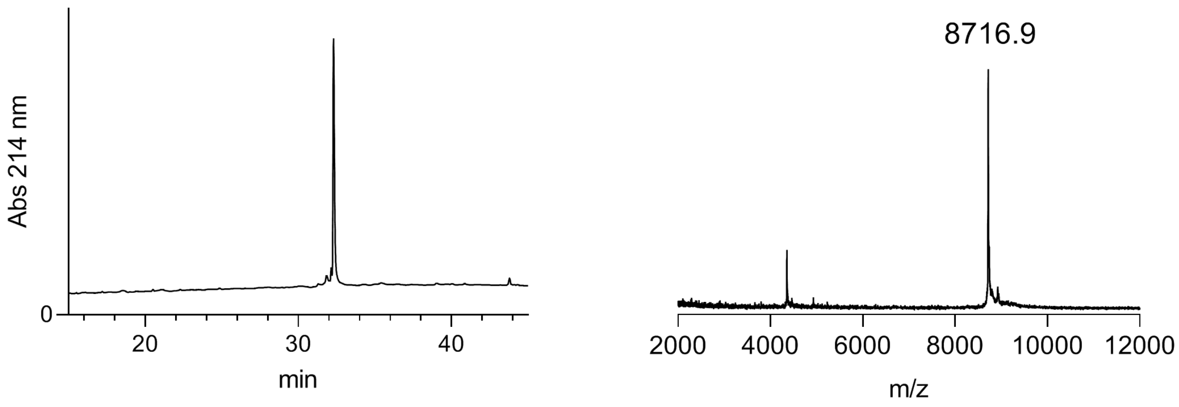


*P*[SYN] *Na*$_v$*1.5_ΔK1500_NM*

The full peptide was assembled according to the general procedure 'C-to-N directed ligations' described above. Preparative HPLC purification followed by lyophilization yielded the peptide as a fluffy solid (1.6 mg as a TFA salt; yield 28%).

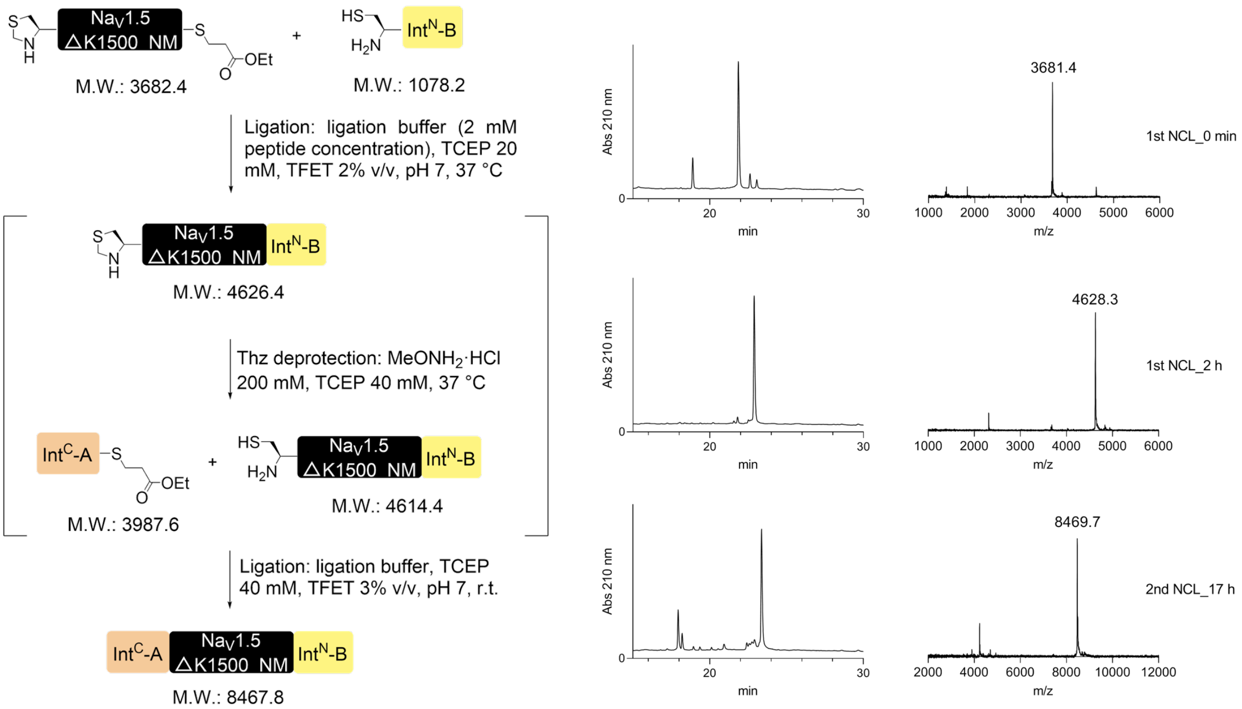

Prep-HPLC purification conditions: 0–10% eluent II in eluent I (5 min gradient) followed by 10–45% eluent II in eluent I (45 min gradient).

Low resolution MS (MALDI-TOF): calc. $[C_{375}H_{611}N_{102}O_{112}S_4]^+$ $[M + H]^+$: 8467.4 Da; found: 8466.1 Da.

HR-MS: calc. $[M + 9H]^{9+}$: 941.7204 Da (941.8318 Da); found: 941.8309 Da (941.7196 Da)

calc. $[M + 10H]^{10+}$: 847.6490 Da (847.7493 Da); found: 847.7489 Da (847.6487 Da)

calc. $[M + 11H]^{11+}$: 770.6816 Da (770.7727 Da); found: 770.7723 Da (770.6812 Da)

calc. $[M + 12H]^{12+}$: 706.5421 Da (706.6256 Da); found: 706.6256 Da (706.5420 Da)

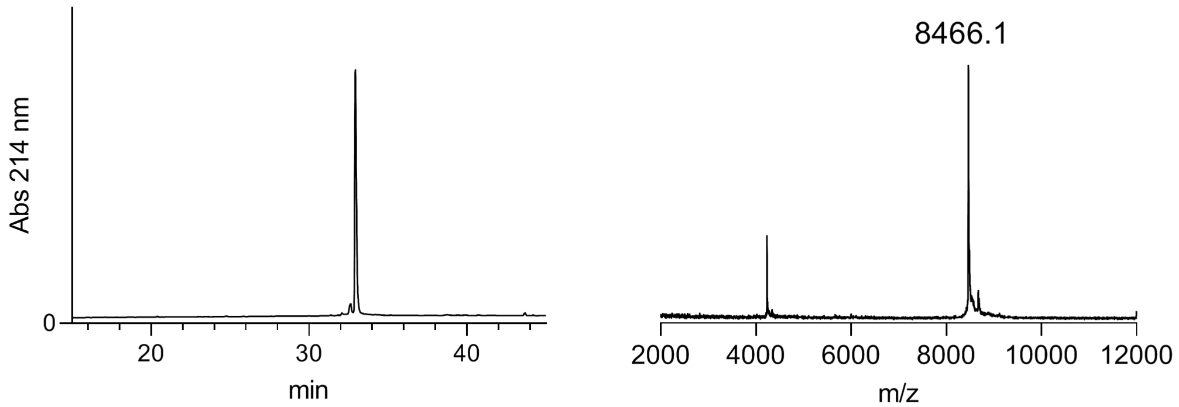


*$P^{SYN}{}_{Na_v1.5\_\Delta K1500\_phY1495}$*

The full peptide was assembled according to the general procedure ‘C-to-N directed ligations’ described above. Preparative HPLC purification followed by lyophilization yielded the peptide as a fluffy solid (1.7 mg as a TFA salt; yield 30%).

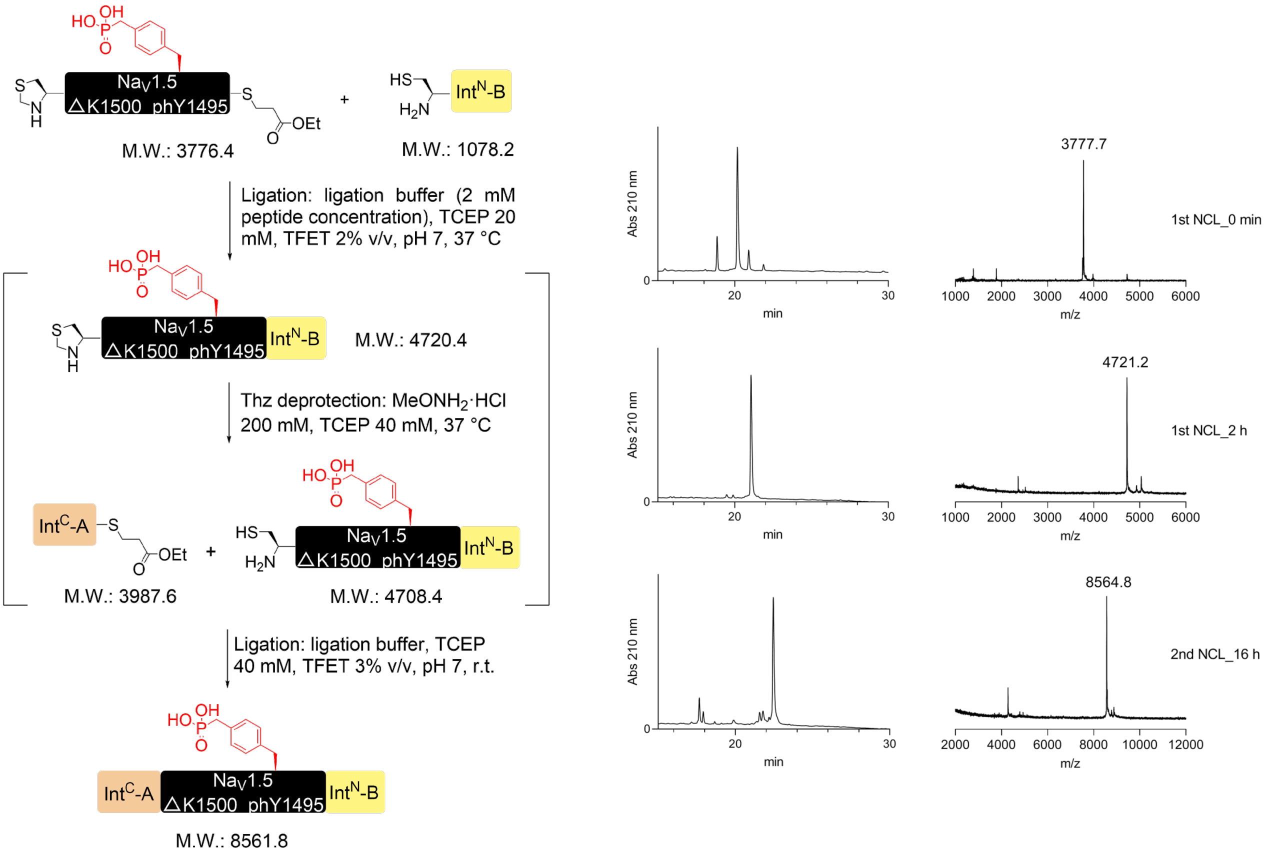


Prep-HPLC purification conditions: 0–10% eluent II in eluent I (5 min gradient) followed by 10–45% eluent II in eluent I (45 min gradient).

Low resolution MS (MALDI-TOF): calc. $[C_{376}H_{614}N_{102}O_{115}PS_4]^+$ $[M + H]^+$: 8560.4 Da; found:  8559.4 Da.

HR-MS: calc. $[M + 9H]^{9+}$: 952.1628 Da (952.2742 Da}; found: 952.2758 Da (952.1646 Da)

calc. $[M + 10H]^{10+}$: 857.0472 Da (857.1475 Da); found: 857.1483 Da (857.0483 Da)

calc. $[M + 11H]^{11+}$: 779.2254 Da (779.3166 Da); found: 779.3169 Da (779.2259 Da)

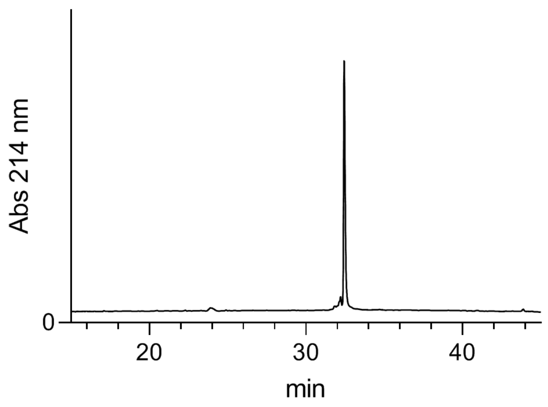
Abs 214 nm
0
20
30
40
min

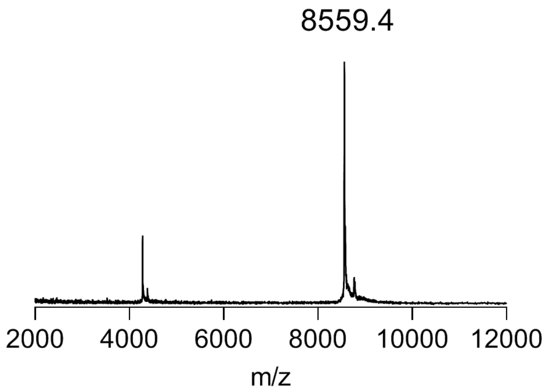
8559.4
2000
4000
6000
8000
10000
12000
m/z
